# Biosurfactant production maintains viability in anoxic conditions by depolarizing the membrane in *Bacillus subtilis*

**DOI:** 10.1101/720532

**Authors:** Heidi A. Arjes, Lam Vo, Caroline Marie Dunn, Lisa Willis, Christopher A. DeRosa, Cassandra L. Fraser, Daniel B. Kearns, Kerwyn Casey Huang

## Abstract

The presence or absence of oxygen in the environment is a strong effector of cellular metabolism and physiology. Like many eukaryotes and some bacteria, *Bacillus subtilis* is an obligate aerobe that primarily utilizes oxygen during respiration to generate ATP. Despite the importance of oxygen for *B. subtilis* survival, we know little about how oxygen is consumed during growth and how populations respond to shifts in oxygen availability. Here, we find that when oxygen was depleted from stationary phase cultures ∼90% of *B. subtilis* 3610 cells died and lysed due to autolysin activity; the remaining cells maintained colony-forming ability. Interestingly, the domesticated 168 strain maintained a higher optical density than 3610 during oxygen depletion due to the formation of cell-wall-less protoplasts, but the remaining, rod-shaped cells were >100-fold less viable than 3610. We discovered that the higher viability in 3610 was due to its ability to produce the antibacterial compound surfactin, as surfactin addition rescued 168 viability and also increased yield in aerobic growth. We further demonstrate that surfactin strongly depolarizes the *B. subtilis* membrane, and that other known membrane-potential disruptors restore viability to 168. These findings highlight the importance of surfactin for survival during oxygen-depleted conditions and demonstrate that antimicrobials normally considered harmful can instead benefit cells in stressful conditions when the terminal electron acceptor in respiration is limiting.

## Introduction

Many species across all domains of life use oxygen as the terminal electron acceptor during aerobic respiration, which produces approximately ten-fold more ATP per glucose molecule than via glycolysis and fermentation in the absence of respiration. In most human cells, oxygen depletion causes exhaustion of ATP and eventual death, either through lysis caused by osmotic stress due to the inability to regulate osmolyte levels [1, 2] or through activation of signaling cascades that lead to apoptosis [3]. Thus, maintaining concentration gradients of ions across cell membranes is of paramount importance when oxygen is lacking. Like human cells, certain microbes such as the pathogen *Mycobacterium tuberculosis* as well as most fungi (with the exception of yeasts) [4] are considered strict aerobes due to their inability to make ATP in the absence of oxygen. Under conditions of rapidly depleted oxygen, *M. tuberculosis* cells show a loss in viability [5, 6]. The related, soil-dwelling species *Mycobacterium smegmatis* also loses viability upon oxygen depletion, and the remaining viable cells maintain the redox balance using hydrogen fermentation and activate stress response genes critical for survival [7, 8]. Despite the clinical importance of *M. tuberculosis*, surprisingly few studies have interrogated how these and other strict aerobes respond to oxygen depletion and the genes responsible for survival.

The Gram-positive model bacterium *Bacillus subtilis* is considered a strict aerobe when grown in the absence of nitrate or nitrite [9]. *B. subtilis* is naturally found in the soil and is used as an additive to help prevent infections and promote growth in plants [10, 11]. In the soil, *B. subtilis* undergoes constant shifts in oxygen concentration, as oxygen is readily available in dry soils but diffuses less and becomes depleted in wet or flooded soils following a rain [12]. Early observations of *B. subtilis* culture lysis upon a shift to anoxic environments have yet to be further characterized [13], and it remains a mystery whether

*B. subtilis* has strategies to cope with oxygen limitation. *B. subtilis* has long been domesticated in the lab, leading to a multitude of genetic tools, strain libraries, and online databases and resources [14–16]. The high level of genetic relatedness between biofilm-forming “wild” strains and derivative non-biofilm-forming laboratory strains has been exploited to identify genetic differences that underlie biofilm community behaviors [17]. For instance, the commonly studied laboratory strain 168, which is derived from the biofilm-forming strain 3610, lacks an extrachromosomal plasmid and harbors several point mutations that reduce or abolish social behaviors such as matrix production [18]. Notably, 3610 produces the small molecule surfactin, a powerful surfactant that has previously been implicated in swarming motility [19–21]. Surfactin has also been shown to kill fungi [22] and some bacteria *in vitro* [22–26]. However, fitness benefits of surfactin production in the context of planktonic cultures have yet to be identified, although some surfactants can accelerate oxygen diffusion through the air-water interface [27].

Here, we characterize the interplay between oxygen availability and surfactin production during the growth and death of *B. subtilis* cultures. We show that oxygen becomes limiting in the culture during the transition to stationary phase and that surfactin secretion improves growth yield in stationary phase by increasing oxygen availability. During a shift to anoxic conditions, we demonstrate that the majority of *B. subtilis* cells die and lyse due to the activity of the LytC autolysin and surfactin. Finally, we discover that surfactin maintains the viability of the remaining cells by causing membrane depolarization that allows these cells to survive until oxygen is restored.

## Results

### Oxygen depletion leads to rapid death and lysis of most *B. subtilis* cells

When culturing the biofilm-forming *B. subtilis* strain 3610 in LB, we noticed that once a test tube containing a late exponential culture was shifted from a shaking incubator to a stationary rack on the bench, the opacity of the tube decreased continuously over a period of 10 hours, suggesting that cells were dying and lysing (Fig. 1A). Consistent with lysis, microscopic observation of cultures left to sit on the bench for 10 hours revealed substantial phase-gray cell remnants in addition to phase-dark rod-shaped cells (Fig. 1A). We conclude that *B. subtilis* 3610 cultures exhibit death and lysis during static incubation after cessation of rapid growth.

**Figure 1:**
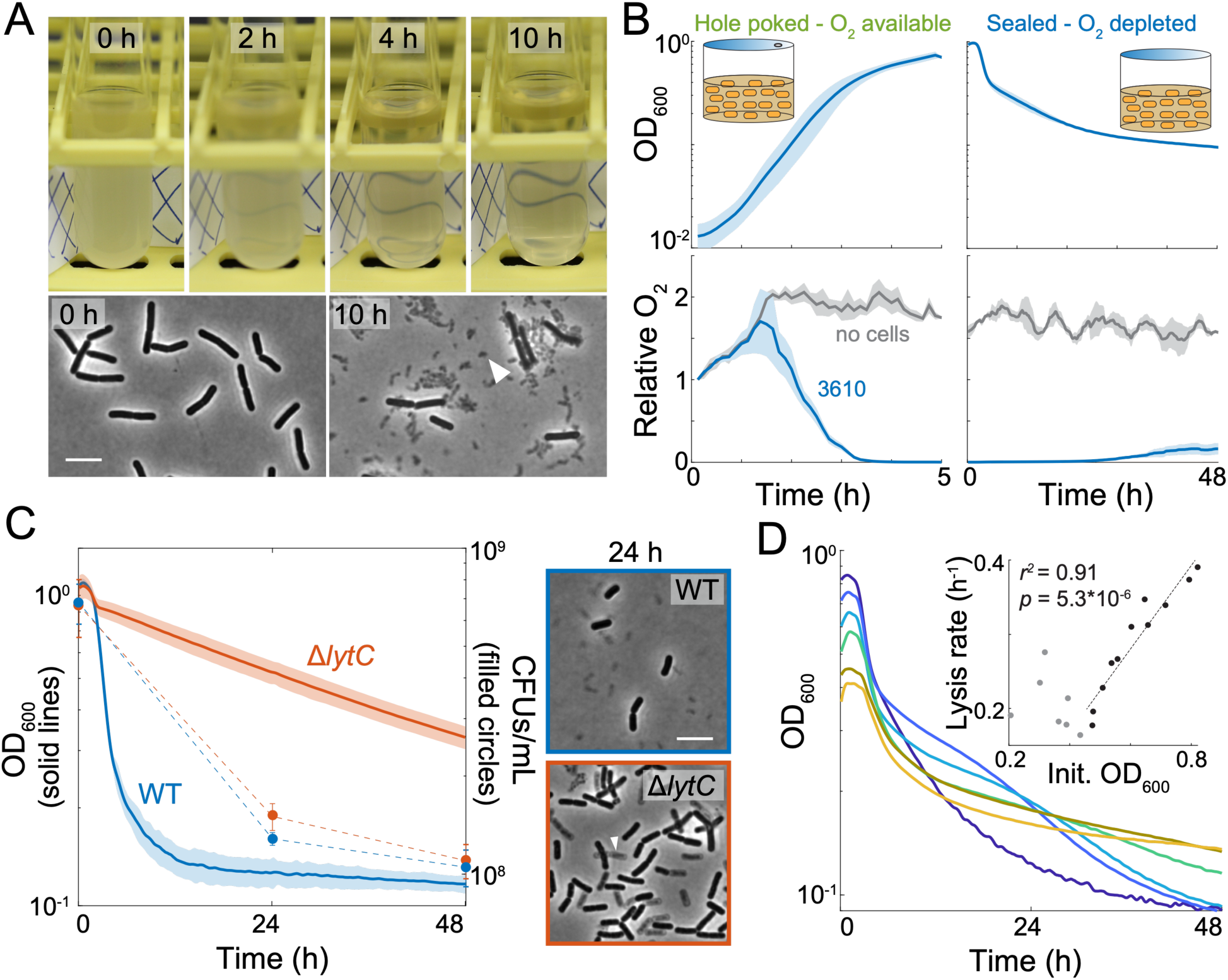
*B. subtilis* 3610 lyses due to oxygen depletion. (A) *B. subtilis* cultures lyse when not shaking. Wild-type strain 3610 (WT) was grown in a test tube and then incubated at room temperature without shaking for 10 h. Phase-contrast images were acquired at 0 and 10 h. Scale bar: 5 μm; arrowhead points to cell debris. (B) Oxygen is depleted during exponential growth and remains low throughout subsequent oxygen depletion while cultures are sealed. Cells were cultured with oxygen-sensitive nanoparticles (Methods). OD_600_ (top) and the relative oxygen levels (bottom, oxygen level at first timepoint normalized to 1) of the cultures were measured. Lines represent the average and shading represents one standard deviation (SD), *n*=3. (C) LytC is necessary for lysis. Left: cultures were grown to an OD_600_∼1 and then oxygen was depleted at *t*=0 h and OD_600_ was monitored. Shading represents 1 SD, *n*=3. Despite the differences in OD, the Δ*lytC* 3610 mutant has similar viability to wild type (*p*=0.07 at 24 h and 0.67 at 48 h, Student’s t-test). Right: phase-contrast images of wild-type and Δ*lytC* cells at 24 h post-oxygen depletion. Scale bar: 5 μm; arrowhead points to phase-gray, lysed cell. (D) Culture lysis is strongly correlated with initial cell density when the initial OD_600_>0.45. Representative lysis curves of 6 cultures that vary in initial OD_600_ (see Fig. S1 for other independent replicates). Inset: the maximum lysis rate vs. initial OD_600_. A linear regression analysis was performed on all data with initial OD_600_>0.45.

The static culture would be limited in oxygen diffusion, and as *B. subtilis* is a strict aerobe that relies on its use of oxygen as a terminal electron acceptor during respiration, we hypothesized that the death in the culture was due to oxygen limitation. To measure oxygen levels during oxygen growth and depletion, we added phosphorescent oxygen-sensitive nanoparticles (Methods) to cell cultures in microtiter plates and measured optical density (OD) and light emission over time. To establish a controlled environment, we grew cultures in sealed 96-well microtiter plates with a hole poked in the seal to allow oxygen exchange, or limited oxygen by completely sealing the wells (Fig. 1B, top); we utilized linear shaking to prevent an oxygen gradient in the culture. In media without cells, oxygen initially diffused into the media due to shaking, and then remained at an approximately constant value (Fig. 1B, bottom). In media with cells, oxygen levels initially increased (again due to shaking) but then began to decrease when the culture reached an OD_600_∼0.05, indicating that the cells were consuming the oxygen faster than it dissolved into the media (Fig. 1B). When cultures reached an OD_600_∼0.5, oxygen levels were undetectable but the culture continued to grow (Fig. 1B), presumably because the cells rapidly consumed any oxygen that dissolved into the media.

When the cultures reached an OD_600_∼0.8, we sealed the wells to abolish oxygen exchange in the headspace, and returned the cultures to a shaking environment. As expected, the measured oxygen levels in the cultures remained low (Fig. 1B, lower right). Within 2 hours, the OD_600_ began to decrease, reminiscent of our observations of OD loss in standing test tubes (Fig. 1A), and plateaued at ∼0.1 after 48 h (Fig. 1B, upper right). Consistent with the OD drop upon sealing the well, cell viability (defined here as the ability of a cell to form a colony on an agar plate) dropped approximately ten-fold after 48 h (Fig. 1C). These observations suggest that the loss of OD in standing tubes and in agitated sealed plates was due to cell lysis during oxygen depletion.

To further explore the correlation between cell lysis and oxygen depletion, we investigated the effect of cell density on hypoxia-induced cell lysis by growing cultures aerobically to different optical densities prior to sealing the wells. Regardless of starting culture density (OD_600_ from 0.4 to 0.8), the OD_600_ either stayed approximately constant or increased slightly for ∼1 hour after sealing, and then the cultures exhibited large decreases in OD over time (Fig. 1D, S1A). The maximum lysis rate (the absolute value of the most negative slope of the ln(OD_600_) curve) increased with increasing starting OD, such that the cultures that began at the highest density (OD600∼0.8) had a lysis rate of 40% decrease per hour (Fig. 1D). Since the lysis behavior varied with initial OD, we standardized all further experiments by growing 3610 cells to an OD_600_ of ∼0.9-1.1 before cutting off oxygen exchange. Transfer of OD∼1 cultures into an anaerobic chamber resulted in an even higher maximum lysis rate of 86% decrease per hour (Fig. S1B). From these data, we infer that populations at high density deplete the remaining oxygen more rapidly, and that rapid oxygen depletion more readily triggers cell lysis.

### A cell wall autolytic enzyme is necessary for lysis of non-viable cells upon oxygen depletion

In addition to losing viability upon oxygen depletion, the non-viable population also lysed and was removed from the population of intact cells, as indicated by the substantial decrease in optical density. Given the importance of the cell wall for maintaining cellular integrity in bacteria, we hypothesized that this lysis involves the activity of cell wall autolytic enzymes. Cell wall growth requires both insertion of new material and cleavage of the existing peptidoglycan by autolysins [28]. In *B. subtilis*, the major autolysin LytC is under control of the vegetative sigma factor **σ**^A^ and the alternative sigma factor **σ**^D^ [29, 30], preventing the unchecked breakdown of the cell wall when insertion is disrupted [31]. We measured the OD of a Δ*lytC* 3610 mutant and found that the mutant exhibited reduced lysis upon oxygen depletion (Fig. 1C). As a specificity control, deletion of another **σ**^D^-dependent autolysin LytD phenocopied wild-type behavior (Fig. S1C), suggesting that LytC is the primary autolysin activated in oxygen-depleted conditions. Despite the higher biomass in the Δ*lytC* cultures after oxygen depletion, a similar proportion of cells (∼10%) retained viability as in wild-type cultures (Fig. 1C). Single-cell imaging of the oxygen-depleted cultures revealed mixed populations of rod-shaped cells, some of which were phase-gray “ghosts” and others that were phase dark (Fig. 1C). Time-lapse microscopy demonstrated that only a portion of the phase-dark rod-shaped Δ*lytC* cells were capable of resuming growth (Fig. S1D, movie S1, S2), consistent with our plating efficiency data (Fig. 1C). These data indicate that lysis and cell viability are genetically separable phenotypes, since many Δ*lytC* mutant cells remain intact yet still lose viability. Thus, since LytC degrades the cell wall but does not impact the viability of 3610 cultures upon oxygen depletion, we conclude that lysis is downstream of viability loss during oxygen depletion.

### Laboratory-domesticated *B. subtilis* strains exhibit more viability loss despite reduced lysis upon oxygen depletion

*B. subtilis* strain 168, a genetic derivative of 3610, is commonly used in research laboratories as it was selected for dispersed growth in culture and improved genetic tractability [17]. In microtiter plate assays, we found that upon oxygen depletion the OD of 168 cultures decreased at a lower rate and remained at a higher level throughout 48 h of monitoring (Fig. 2A). Imaging of oxygen-depleted 168 cultures revealed the presence of round cells in addition to rod-shaped cells (Fig. 2B). Some round cells were intact and did not stain with propidium iodide (PI), while others had compromised membranes and thus stained brightly with PI (Fig. 2B). In cell cultures that were first stained with fluorescent D-amino acids (FDAAs) to label cell walls, the vast majority of the rounded cells did not retain FDAA staining, indicating that they were protoplasts without cell walls (Fig. S2A). Indeed, a 168 Δ*lytC* strain did not form round cells and exhibited less lysis (Fig. S2B), indicating that the protoplasts released in 168 are more fragile than their walled counterparts. Thus, we conclude that LytC activity during oxygen depletion degrades the cell wall in 168, releasing membrane-bound protoplasts that account for the higher OD in 168 cultures compared with 3610 cultures.

**Figure 2:**
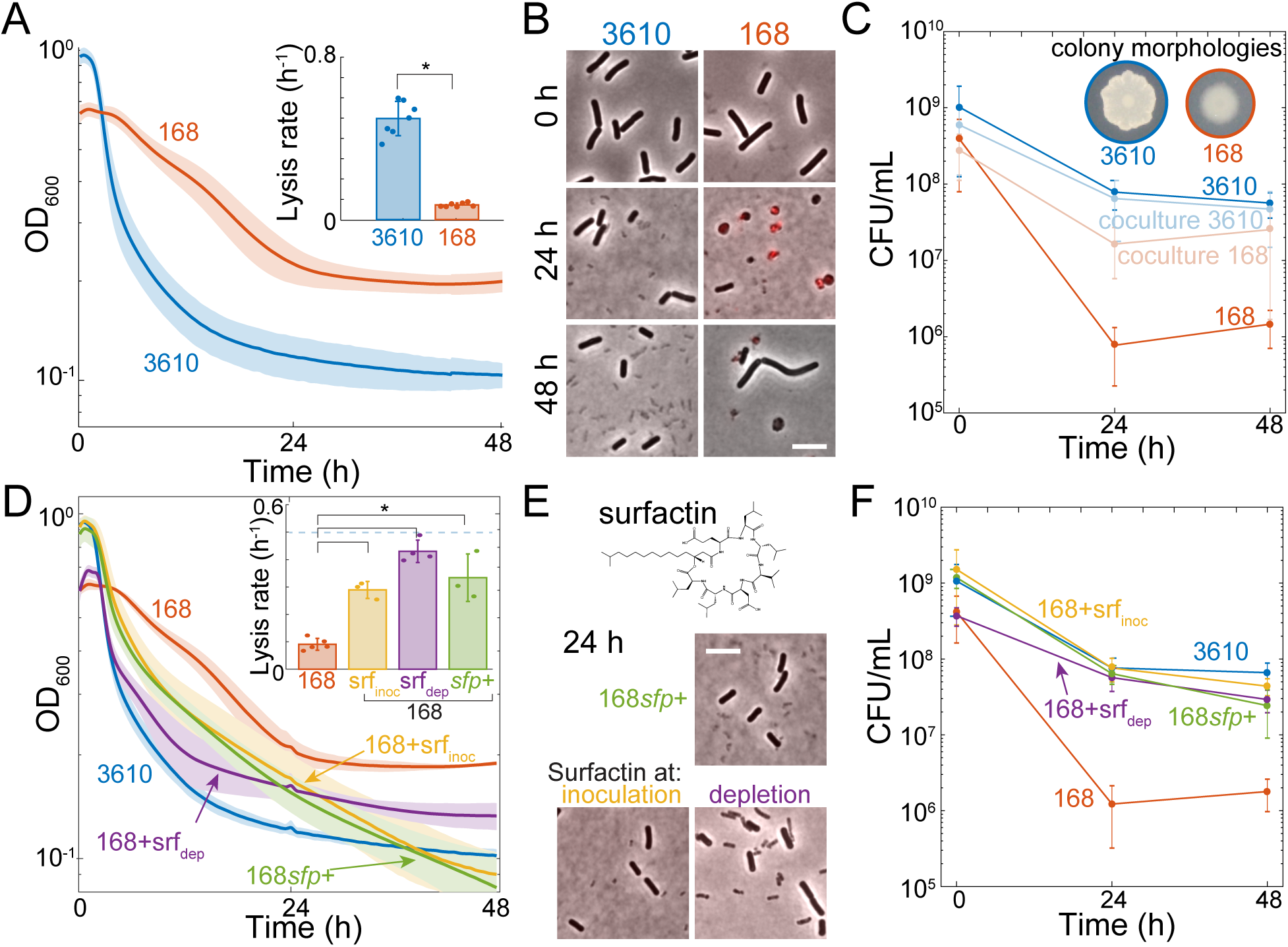
Surfactin production is necessary to maintain viability. (A) *B. subtilis* strain 168 lyses less upon oxygen depletion than 3610*. B. subtilis* 3610 and 168 strains were grown aerobically and then depleted for oxygen at 0 h. Lines represent the average and shading represents 1 SD, *n*=7. Inset: maximum lysis rate of 3610 culture is significantly higher than that of 168 (*: *p*=1.7×10^−8^, Student’s t-test; Methods). Circles show individual experiment values, and error bars represent 1 SD. (B) Many 168 cells form cell-wall-less protoplasts upon oxygen depletion. Merge of phase-contrast and fluorescence images of propidium iodide (PI)-stained cells at 0, 24, and 48 h post oxygen depletion. Red indicates membrane-compromised cells. (C) Co-culturing 168 with 3610 rescues its viability upon oxygen depletion. 168 viability in monoculture is significantly different than 3610 (*: *p*<0.005; student’s t-test). Inset: 3610 and 168 have distinct colony morphologies when plated on LB. Error bars represent 1 SD*, n*=3-5. (D) Culturing with exogenous surfactin increases lysis of 168 cultures upon oxygen depletion. OD curves during oxygen depletion of 3610, 168, and 168 genetically rescued for surfactin production (168*sfp*+) or with 48 µM exogenous surfactin added before growth (srf_inoc_) or at depletion (srf_dep_). Lines represent the average and shading represents 1 SD, *n*=3-5. Inset: maximum lysis rates (*: *p* <0.001; Student’s t-test). (E) Culturing with exogenous surfactin eliminates protoplasts from 168. Top: surfactin molecular structure. Bottom: phase-contrast and PI fluorescence imaging at 24 h post-oxygen depletion of 168*sfp*+ cells or with exogenous surfactin. (F) Surfactin restores the viability of 168 cultures to near 3610 levels upon oxygen depletion. Error bars represent 1 SD, *n* = 3-5. Surfactin-treated 168 (srf_inoc_, srf_dep_, and 168*sfp*+) cultures are each significantly different than 168 (*p*<0.001 at 24 h, *p*<0.01 at 48 h, Student’s t-test). By contrast, the viability of surfactin-treated 168 cultures are not significantly different than that of 3610 at 24 h (*p*>0.2, Student’s t-test).

Despite having a higher OD_600_ than 3610 cultures, the colony-forming units (CFU) in 168 cultures depleted of oxygen for 24 h decreased ∼100-fold relative to 3610 (Fig. 2C). Indeed, time-lapse imaging on fresh medium with oxygen showed that despite having intact cell envelopes, only ∼1% of the rod-shaped 168 cells were able to grow and divide, whereas ∼95% of the 3610 cells exhibited growth (Fig. S2C, Movie S3,S4). We never observed growth either in increased cell number or increased cell size during 12 h time-lapse imaging of any of the protoplasts on fresh LB or on filtered spent medium, suggesting that the protoplasts formed in strain 168 are not viable. We conclude that despite reduced cell lysis, 168 cultures experience a much more drastic loss of viability upon oxygen depletion.

### Surfactin increases lysis and restores viability to the domesticated strain 168

Strains 3610 and 168 differ in a number of chromosomal loci, and 168 is defective in phenotypes related to secretion of extracellular products that support multicellular behaviors such as biofilm formation and swarming motility [18]. To determine whether extracellular products were responsible for the oxygen-related phenotypic differences between the two strains, we subjected a 1:1 volumetric mixture of the strains to oxygen depletion. The viability of 168 (which has a colony morphology visually distinct from that of 3610) increased dramatically in the co-culture (Fig. 4C). A candidate compound that could restore viability to 168 is surfactin, which is a strong surfactant that creates K^+^-permeable pores and solubilizes lipid bilayer vesicles *in vitro* [32]. Surfactin is produced by 3610 but not 168, due to a mutation in *sfp*, which encodes an enzyme necessary to activate the surfactin biosynthesis complex [19]. We found that exogenous surfactin addition at either the time of inoculation of the culture or at the initiation of oxygen depletion increased the lysis rate of 168 and eliminated protoplasts (Fig. 2D,E), consistent with surfactin-induced protoplast lysis.

Remarkably, exogenous surfactin addition restored the viability of oxygen-depleted 168 cultures to similar levels as 3610 (Fig. 2F). Moreover, a 168 strain with *sfp* genetically complemented to restore surfactin production phenocopied the surfactin-treated 168 cultures in terms of lysis, cell morphology, and viability (Fig. 2F, S2). Thus, we conclude that surfactin production both promotes protoplast lysis and maintains viability in oxygen-depleted *B. subtilis* cultures.

### Surfactin improves growth yield of the domesticated strain 168 by increasing oxygen diffusion

One intriguing distinction between the aerobic growth of 3610 and 168 was the divergence of the growth curves around OD_600_∼0.3; after this point, 3610 cultures completed another ∼1.5 mass doublings to reach OD_600_∼0.9, while 168 cultures underwent only 1 mass doubling to reach OD_600_∼0.6 over the same period (Fig. 3A). We noted that a surfactin-supplemented culture experienced more growth after the OD reached ∼0.3 than untreated cultures, resulting in a higher overall yield similar to that of 3610 (Fig. 3B). When surfactin was added just prior to the divergence in growth curves, the yield was similarly restored (Fig. 3B). Thus, we inferred that surfactin production was responsible for the increase in yield of 3610 relative to 168. The detergent Tween 80 similarly increased growth yield in 168 cultures (Fig. 3C), indicating that the detergent properties of surfactin were likely responsible for the increased yield.

**Figure 3:**
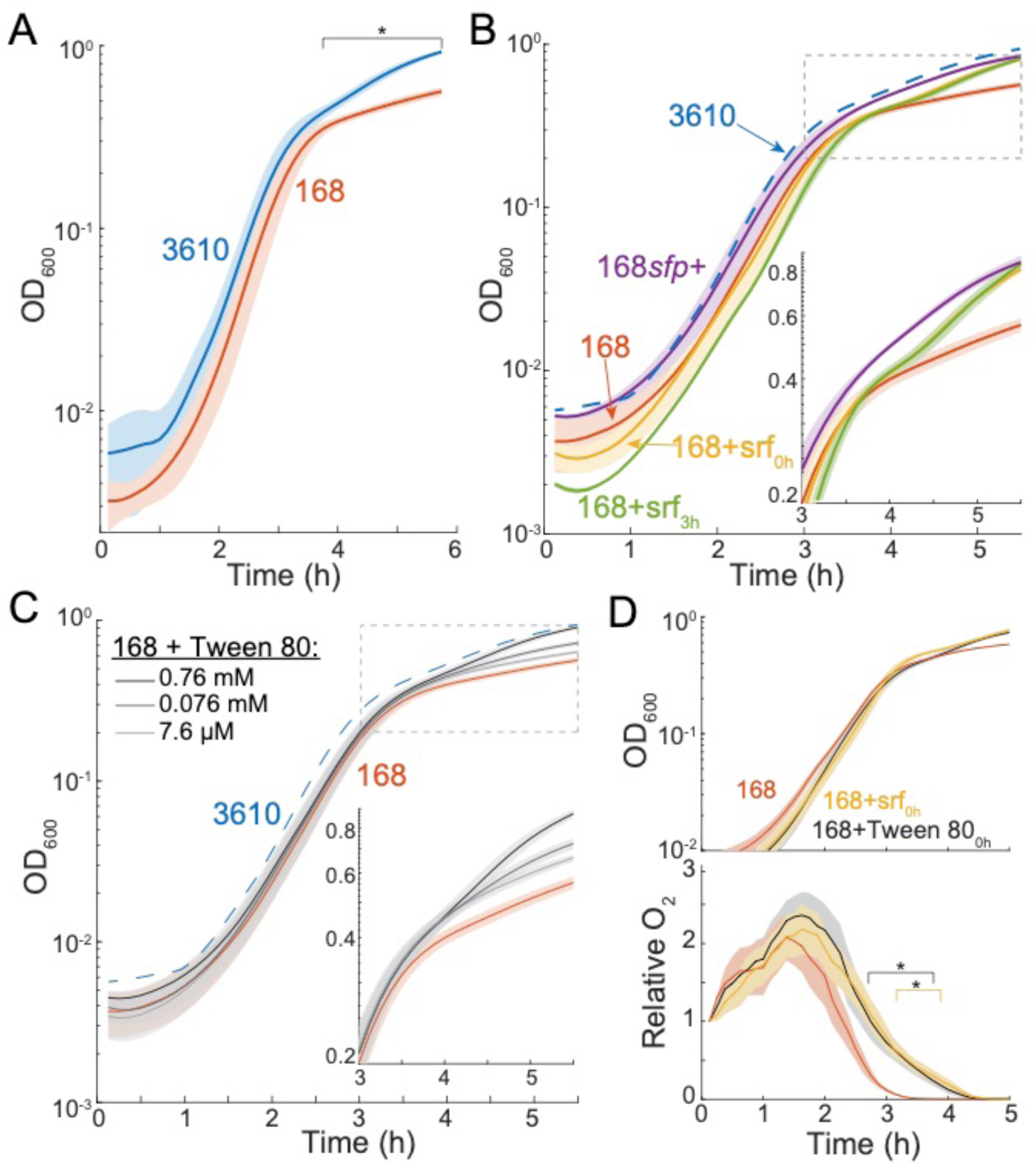
Surfactin restores the growth yield of 168 due to its detergent properties. (A) 3610 aerobic cultures achieve a higher growth yield than 168. Lines represent the average and shading represents 1 SD, *n*=3. *: time period over which 168 growth differed significantly from that of 3610 (*p<*0.05, Student’s t-test). (B) Surfactin addition rescues growth yield. Growth curves of 168 with surfactin restored genetically (168*sfp*+) or 48 μM added exogenously at inoculation (168+srf_0h_) or at t = 3 h (168+srf_3h_). Lines represent the average and shading represents 1 SD, *n*=3. The mean 3610 growth curve from (A) is shown as dotted blue line. Inset: zoom-in to the period of growth divergence. (C) Tween 80 addition rescues growth yield in a concentration-dependent manner. Lines represent the average and shading represents 1 SD, *n*=3. The mean 3610 growth curve from (A) is shown as dotted blue line. Inset: zoom-in to the period of growth divergence. (D) Surfactin addition increases oxygen levels during late exponential phase. Relative oxygen levels (bottom) during growth (top) of 168 with surfactin (48 μM) or Tween 80 (0.76 mM) added. *: time period over which oxygen levels of 168+surfactin (yellow) or 168+Tween 80 (black) were significantly different from 168 cultures (*p*<0.05, Student’s t-test).

Certain compounds have the ability to increase oxygen diffusion in liquid [33], and it has been proposed that oxygen diffusivity is rate-limiting for the function of some biological systems [34]. To test whether surfactin increased oxygen in the media, we used our oxygen nanoprobe to compare oxygen levels in 168 cultures with and without added surfactin. We found that exogenous surfactin increased the oxygen levels in cultures during late-exponential phase, when oxygen would normally be depleted lower than our limit of detection (Fig. 3D). A similar increase occurred due to addition of Tween 80 (Fig. 3D). A biophysical model incorporating diffusion and cellular consumption of oxygen predicted that the observed ∼1.2-fold increase in peak oxygen level is consistent with a ∼1.4-fold increase in diffusion rate (Methods). Thus, we infer that the presence of surfactin results in higher growth yield due to the enhanced availability of oxygen during the transition to stationary phase when oxygen would otherwise be limiting for growth.

### A transposon screen supports surfactin production as the main determinant of survival upon oxygen depletion

Given the large decreases in viability after 24 h of oxygen depletion, we performed a genetic screen to attempt to identify any mutations that would increase the survival of 168 and 3610 without oxygen. We made 20-30 independent transposon libraries of 5000-10,000 individual transposon mutants per library in each background and subjected these libraries to oxygen depletion, hypothesizing that any mutants with enhanced survival would be enriched (Fig. S3A). We could not identify any such 168 mutants, suggesting that there is not an easily obtainable loss-of-function mutant in a surfactin-independent pathway to increase viability upon oxygen depletion.

We identified two categories of mutants in the 3610 background with decreased lysis upon oxygen depletion: surfactin production (*comA*, *comP*, *srfAA*, and a hit upstream of *rghR* (*rghR*_us_)) and flagella-related (*fliI, fliJ*, and *fliF*) (Fig. S3B, Table S1). *srfAA* encodes the surfactin synthetase subunit A, which is part of the enzyme that synthesizes surfactin (Fig. 4A) [19, 35]. ComA and ComP form a two-component system that activates surfactin production at high cell density [36]. RghR regulates RapG and RapH, two repressors of the *srfA* operon (Fig 4A) [37]. The flagella-related mutants disrupted *fliI* and *fliJ*, which encode accessories of the flagellar type III secretion apparatus, and *fliF*, which encodes the flagellar basal ring [38–41]. In-frame markerless deletions of these mutants had ∼4-fold lower viability than the parent after 24 h of oxygen depletion (Fig. S3C), which we hypothesized was due to the surfactin in the pooled library cultures increasing their survival advantage in the mixed population but not in isolation. Indeed, upon exogenous surfactin addition, all mutants responded with increased lysis and removal of protoplasts/cell debris (Fig. 4B,C, S3D). Taken together, the fact that we obtained no hits that increased viability in a 168 background and that all hits in a 3610 background are directly related to surfactin production or respond to surfactin points to surfactin as the primary determinant of viability in the absence of oxygen.

**Figure 4:**
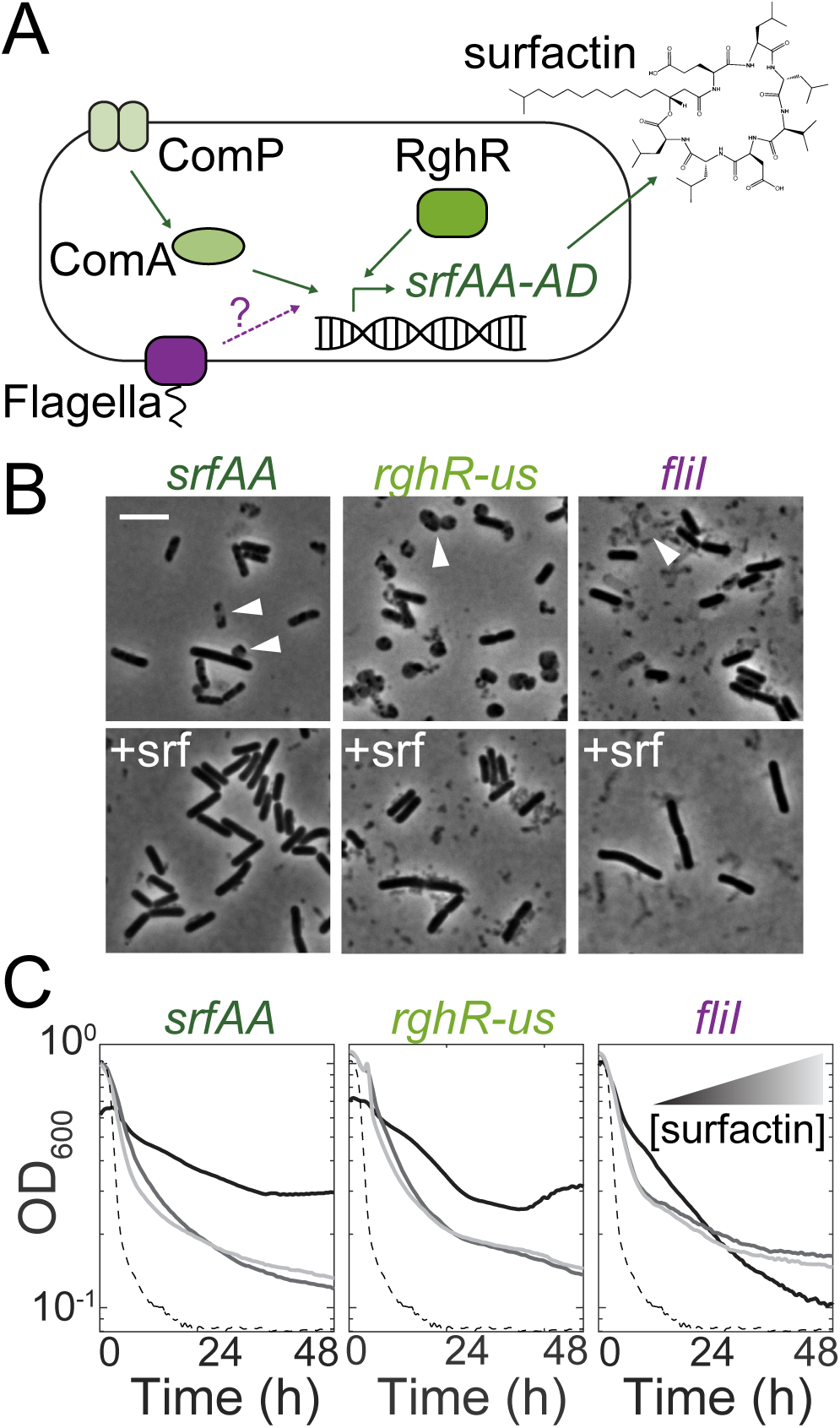
Transposon mutagenesis identifies genes that impact lysis during oxygen depletion. (A) Schematic of regulation of the surfactin synthetase gene operon (*srfAA-AD*). Known regulators of SrfAA are shown in green. Our data suggests flagellar proteins (purple) may also regulate surfactin. (B) Surfactin-treated cultures of the transposon-disrupted mutants have fewer protoplasts and cell debris. Phase-contrast images of mutants depleted of oxygen for 24 h with and without 48 μM exogenous surfactin. Top: arrowheads show protoplasts, phase-gray dead cells, and cell debris, all of which were not observed in the surfactin-treated cultures. (C) Transposon hits exhibit faster lysis when treated with exogenous surfactin. Black curves are without surfactin, medium and light gray curves are with 24 μM and 48 μM surfactin, respectively. The dashed line is the parent (3610).

### Surfactin restores viability by depolarizing the membrane

*In vitro*, surfactin creates potassium ion-permeable pores in lipid bilayers [32]. If this behavior occurs *in vivo*, such pores would reduce the strength of the potassium ion gradient across the cell membrane and alter membrane potential. Thus, we tested the energetic state of cells after various chemical treatments using the membrane potential-sensitive dyes DiSC_3_(5) and ThT [42, 43]. DiSC_3_(5) is taken up by cells and the fluorescence signal is initially quenched. Agents that depolarize cells release DiSC_3_(5) into the medium, resulting in an increased signal [42]. By contrast, ThT enters cells with polarized membranes and fluoresces inside the cell; upon depolarization, ThT exits the cells and the fluorescence is reduced [43]. As expected, we found that treatment of 168 cells with valinomycin, a known depolarizing agent that functions as a potassium-specific transporter [44], led to an increase in DiSC_3_(5) and a decrease in ThT signal (Fig. 5A). CCCP, a proton ionophore that dissipates the proton motive force [45], strongly reduced the ThT signal and slightly increased DiSC_3_(5) fluorescence (Fig. S4). Surfactin treatment led to a large increase in DiSC_3_(5) fluorescence and a greater decrease in ThT fluorescence than valinomycin (Fig. 5A), demonstrating that surfactin can strongly depolarize *B. subtilis* cells.

**Figure 5:**
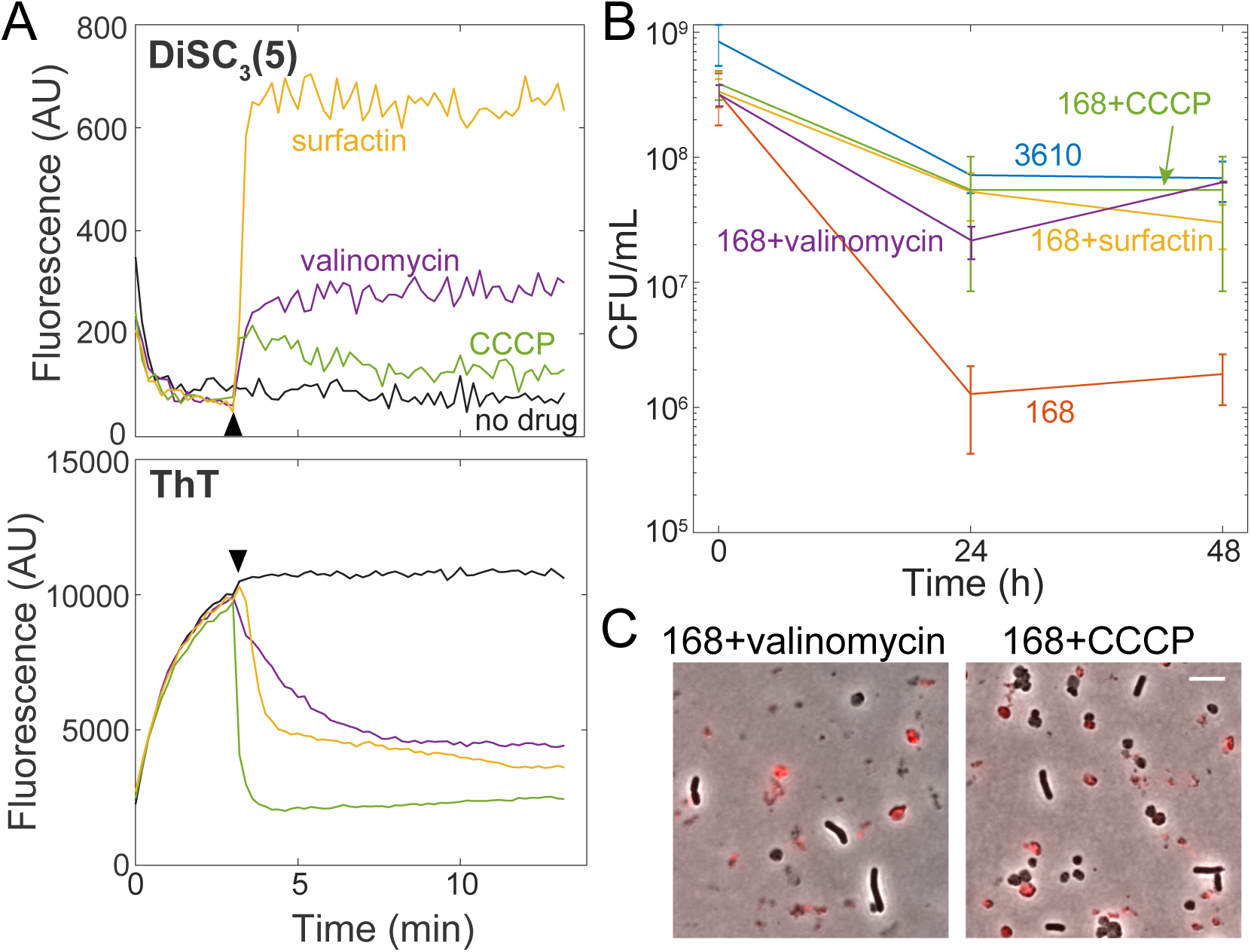
Surfactin maintains viability upon oxygen depletion by depolarizing the membrane. (A) Surfactin depolarizes the membrane in *B. subtilis*. Membrane potential assays of 168 cells using the dyes DiSC_3_(5) (top) and ThT (bottom). The time of addition of surfactin (48 μM), valinomycin (50 μM) and CCCP (5 μM) is marked by the black arrowhead. One representative experimental replicate is shown (other replicates are in Fig. S4). (B) Treatment with the membrane depolarizing agents valinomycin (5 µM) and CCCP (5 µM) restore plating efficiency of *B. subtilis* 168 after oxygen depletion, similar to surfactin (48 µM). Error bars represent 1 SD, *n*=3-5. 168 plating efficiency data are significantly different than those of 168+surfactin, 168+valinomycin, 168+CCCP, and 3610 (*p*<0.005, Student’s t-test). (C) Valinomycin and CCCP-treated 168 cultures exhibit protoplasts, demonstrating that protoplast removal is not necessary for viability enhancement. Overlays of phase-contrast and PI fluorescence (red) images at 24 h post-oxygen depletion. Scale bar: 5 μm.

Based on these findings, we hypothesized that de-energizing the membrane would be sufficient to rescue the colony-forming ability of 168 cultures upon oxygen depletion. Indeed, valinomycin and CCCP both restored viability to a similar extent as surfactin (Fig. 5B). Interestingly, valinomycin- and CCCP-treated cultures had protoplasts after 24 h of oxygen depletion, consistent with surfactin being necessary for protoplast lysis and indicating that protoplast formation is independent from viability maintenance. Taken together, these data indicate that membrane potential dictates the ability of *B. subtilis* cells to survive oxygen depletion.

## Discussion

While surfactin has been recognized to promote swarming motility and complex colony architecture in *B. subtilis* communities such as biofilms [18, 46], its role in planktonic cultures has remained mysterious. Here, we demonstrate three independent roles of surfactin in planktonic cultures. During aerobic growth, the detergent properties of surfactin increase growth yield by increasing the oxygen available to cells entering stationary phase (Fig. 3). During oxygen depletion, surfactin plays two independent roles: [1] it works in tandem with LytC to remove non-viable cells from the culture (Fig. 2), and (2) it depolarizes the remaining cells and thereby maintains their viability during oxygen depletion (Fig. 5).

The increase in growth yield due to surfactin suggests that oxygen is limiting even in aerobic cultures and that increasing oxygen availability can increase yield. This finding has industrial implications as *B. subtilis* is commonly used to produce many biological compounds such as enzymes and antibiotics [47], and maintaining oxygen levels are critical to optimize production of the desired compound [48, 49]. If this oxygen-related growth yield enhancement holds across species, detergent addition may increase bioproduction in other commonly utilized organisms such as *Streptomycetes* species, which are used to isolate numerous antibiotics, and the industrial powerhouse *Escherichia coli*, which is widely used to synthesize many biopharmaceuticals such as insulin [47]. In addition to oxygen, surfactin may also facilitate growth by increasing nutrient diffusion to cells during late exponential phase when nutrients become limiting.

We also found that surfactin has at least two functions during oxygen starvation that are distinct from its role during aerobic growth. Surfactin causes lysis in cells that experience LytC-mediated cell wall degradation (Fig. S2), presumably through membrane disruption. Moreover, surfactin maintains viability of the remaining intact cells through membrane depolarization, which may allow these cells to enter a metabolically inactive state where they can ride out the stress of the oxygen depletion. While it was formally possible that cell lysis helped maintain viability of the remaining cells, the observations that Δ*lytC* mutants have the same viability as wild-type cultures even though the non-viable cells remain intact (Fig. 1C) and that CCCP or valinomycin treatment of 168 cultures rescues viability without removing protoplasts (Fig. 5C) demonstrate that surfactin maintains viability by acting directly upon the membrane potential of the surviving cells.

Generally, OD is used as a proxy for cell number, which holds true for our measurements of 3610 cultures undergoing oxygen depletion as the 10-fold drop in OD measured reflects cell lysis and quantitatively mirrors the decrease in viability (Fig. 1C). However, our work highlights instances during oxygen depletion when biomass measured by OD is uncoupled from viability: both Δ*lytC* 3610 cells and 168 cells remain intact and the cultures have a relatively high OD compared with wild-type 3610, but most cells cannot form colonies (Fig. 2C, S1C). This uncoupling between OD and viability has been observed previously in cell-cycle mutants where cells remain intact and metabolically active but cannot divide and form colonies [51], and may be more prevalent than is currently appreciated, motivating future studies that rely on OD to perform assays to verify culture viability.

Since cell depolarization can actually improve the viability of cells undergoing oxygen depletion, under conditions with limiting terminal electron acceptors, certain antibacterial treatments that inhibit growth may actually keep cells viable. Such a possibility needs to be taken into consideration when treating bacterial infections and removing bacteria in low-oxygen clinical and industrial settings, particularly those in which the microbe primarily gains energy through aerobic respiration and hence the terminal electron acceptor may be limiting. Moreover, it remains generally unclear which antimicrobial treatments will disrupt membrane potential as a side effect of their primary activity or mechanistically how antibiotic treatments affect cells in low oxygen environments, motivating further studies of membrane energetics during growth-inhibition.

The cell-wall breakdown of *B. subtilis* cells during oxygen depletion provides further support for a recently discovered regulatory role of membrane potential in cell-wall synthesis [52]. Indeed, LytC is activated upon sodium azide treatment that deprotonates the cell wall [53, 54] and the activity of peptidoglycan synthesis enzymes in *E. coli* are regulated by pH [55, 56]. In addition, the recent observation that actively growing and dormant cells in a *B. subtilis* culture respond oppositely to an electrical pulse wherein they either hyperpolarize or depolarize, respectively [57], suggests that membrane energetics may explain the observed population heterogeneity in our oxygen-depleted 3610 cultures. Thus, as membrane potential is of utmost importance in metabolism and cell growth, is becoming more appreciated in regulating cell-wall remodeling, and likely feeds back into many additional aspects of cell physiology, bacteria must employ strategies to maintain and/or modulate membrane potential during changing environmental conditions. The ability of surfactin to both alter membrane energetics to maintain viability during oxygen-depletion and to enhance growth in oxygen-limited conditions has likely provided multiple fitness advantages to *B. subtilis* in spatially structured and complex environments such as native soil communities, and strategies for regulating membrane potential may be an important factor for survival of other strict aerobes as well.

## Methods

### Media and growth conditions

All strains and their genotypes are listed in Table S2. Strains were grown in LB (Lennox broth with 10 g/L tryptone, 5 g/L NaCl, and 5 g/L yeast extract). Antibiotics for selection of mutant strains were used as follows: kanamycin (5 μg/mL), MLS (a combination of erythromycin at 0.5 μg/mL and lincomycin at 12.5 μg/mL), chloramphenicol (5 μg/mL), and spectinomycin (100 μg/mL). Surfactin was added at a final concentration of 0.05 mg/mL unless otherwise noted. Strains were cultured either in 5 mL of medium in a test tube on a roller drum or in 200 μL of medium in a 96-well plate in a Biotek Epoch2 spectrophotometer under linear shaking (shaking was set to 567 cycles per minute (cpm), 3-mm magnitude of shaking). For all experiments, the initial inoculum was from a fresh colony struck from a −80 °C freezer stock onto LB 1.5% agar plates and incubated overnight at 37 °C.

### Strain construction

Strains were constructed using SPP1 phage transduction [58]. The donor strain was grown for >6 h in TY medium (LB supplemented with 0.01 M MgSO_4_ and 0.1 mM MnSO_4_ after autoclaving). Ten-fold dilutions of SPP1 phage were added to the culture and 3 mL TY soft (0.5%) agar was mixed with the culture/phage mixture and poured over a TY plate (1.5% agar) overnight. A plate was chosen that exhibited nearly total clearing of cells without a large number of phage-resistant mutants. Five milliliters of TY medium were added to this plate and a 1-mL filter tip was used to scrape up the soft agar. This soft agar/liquid mix was filtered through a 0.4-µm filter. The phage was added to a stationary-phase (grown for 6-10 h culture in TY medium of the recipient strain (10 μL undiluted phage + 100 μL recipient cells, and optionally 900 μL TY medium) and incubated at 37 °C for 30 min, then plated onto LB + antibiotic and 0.01 M sodium citrate (sodium citrate was omitted for MLS selection). Plates were incubated for 24 h and transductants were struck for single colonies to eliminate the phage.

To generate the Δ*lytC* in-frame marker-less deletion construct, the region upstream of *lytC* was PCR-amplified using the primer pair 1427/1428 and digested with SalI and EagI, and the region downstream of *lytC* was PCR-amplified using the primer pair 1425/1426 and digested with EagI and BamHI. The two fragments were then simultaneously ligated into the SalI and BamHI sites of pMiniMAD, which carries a temperature-sensitive origin of replication and an erythromycin resistance cassette [59], to generate pDP299.

To generate the Δ*lytD* in-frame marker-less deletion construct, the region upstream of *lytD* was PCR-amplified using the primer pair 1429/1430 and digested with SalI and EagI, and the region downstream of *lytD* was PCR-amplified using the primer pair 1431/1432 and digested with EagI and BamHI. The two fragments were then simultaneously ligated into the SalI and BamHI sites of pMiniMAD to generate pDP300.

Deletion plasmids were introduced into strain DK1042 (Table S2) via single cross-over integration by transformation at the restrictive temperature for plasmid replication (37 °C) using MLS resistance as a selection. To evict the plasmid, the strain was incubated in 3 mL LB broth at a permissive temperature for plasmid replication (22 °C) for 14 h, and serially diluted and plated on LB agar at 37 °C. Individual colonies were patched on LB plates and LB plates containing MLS to identify MLS-sensitive colonies that had evicted the plasmid. Chromosomal DNA from colonies that had evicted the plasmid was purified and screened by PCR using primers 1427/1426 or 1430/1431 to determine isolates that had retained the Δ*lytC* or Δ*lytD* allele, respectively [59, 60].

In-frame deletions of MLS-knockout mutants were constructed as outlined previously [14]. Briefly, the strain of interest was transformed with pDR244 at 30 °C. Several transformants were struck onto LB at 42 °C. Resultant colonies were patched on MLS and spectinomycin plates to confirm that the plasmid and MLS cassette were lost. Colonies were screened for altered oxygen phenotype.

### Growth and lysis assays

Strains were struck out for single colonies on the evening prior to the experiment. Colonies were inoculated into fresh LB and grown aerobically at 37 °C, either in 5 mL LB on a roller drum or in 200 μL in a Greiner 96-well plate. To deplete oxygen from cultures grown in test tubes, 200 μL were aliquoted into a 96-well plate and the plate was sealed with optical film (Excel Scientific AeraSeal). To grow cultures in the plate reader, 200 μL of stationary-phase inocula were aliquoted into each well of a 96-well plate, with the edge wells containing medium only. Optical film was used to cover the plates, and one hole per well was poked using a 20-gauge needle to allow for air exchange. 168 and 3610 cultures were then grown until the 3610 culture reached an OD_600_∼1.0. To deplete the oxygen, the plate was taken out of the plate reader, the optical seal with holes was removed, and a new seal was placed onto the plate to fully cut off oxygen exchange. We grew and depleted cultures at 37 °C using linear shaking (567 cpm, 3-mm magnitude) and read OD_600_ every 7.5 min in a Biotek Epoch2 spectrophotometer.

### Mariner transposon library construction

The mariner transposon was used to create a library of insertion mutants. The parent strains (HA1235 and HA1414) were struck for single colonies onto MLS plates at 30 °C. One colony per library was grown in 3 mL LB+kanamycin at room temperature overnight. Transposon-insertion libraries were selected by plating 10-fold dilutions of the cultures on prewarmed LB+kanamycin plates and incubating overnight at 37 °C.

### Screen to enhance for mutants that survive better without oxygen

Each library (∼5000-10,000 colonies per library) was inoculated into 5 mL LB and grown on a roller drum at 37 °C to an OD_600_∼1.0. Libraries were then aliquoted into wells of a 96-well plate (200 μL per well) and plates were sealed to deplete oxygen. Oxygen was depleted at 37 °C for 2-8 days for 168 libraries or at 30 °C for 4-8 days for 3610 libraries. At various time points following the start of oxygen depletion, one well of the library was harvested and struck out for single colonies. One or two colonies per well were tested for either enhanced lysis upon oxygen depletion (3610 background) or enhanced colony-forming ability (168 background). Any mutants found to have these phenotypes were back-crossed into the parent using SPP1 phage transduction to ensure the transposon mutation was the causative agent. To map the mutation, genomic DNA was prepped from each mutant. Inverse PCR was carried out using Phusion polymerase with the primers IPCR1 and IPCR2. The PCR products were gel-purified and sequenced using the IPCR2 primer. The sequences were mapped onto the *B. subtilis* genome using BLASTN.

### Oxygen nanoprobe measurements

Relative oxygen levels were measured using the oxygen-sensitive nanoprobe (BF_2_nbm(I)PLA) that emits an oxygen-dependent phosphorescence reading and an oxygen-independent fluorescence reading, which together can be used to calculate the relative oxygen level of the medium [61, 62]. Briefly, the nanoprobe was added at 5% (10 μL into 190 μL) to the inoculum before growth of the cultures. In addition to OD_600_ readings, fluorescence readings (ex/em: 414/450 nm) and phosphorescence readings (ex/em: 415/560 nm, with a 2-ms delay between excitation and emission) were taken using a Biotek Synergy H1 spectrophotometer. Readings were taken every 7.5 min, with incubation at 37 °C and linear shaking (567 cpm, 3-mm magnitude of shaking).

### PI staining and phase microscopy

Five hundred nanoliters of cultures were spotted onto LB pads made with 1.5% agar and 10 μM PI. Once dry, a coverslip was added and cells were imaged on a Nikon Ti-E inverted microscope using a 100X objective (NA: 1.4). Phase and fluorescence (mCherry filter, ex/em: 570/645 nm) images were acquired. Images were processed identically in Adobe Photoshop and merged using FIJI.

### Time-lapse microscopy of oxygen-depleted cultures

The bottom of a rectangular Singer PlusPlate culture plate was used to make a large pad [63], in which 35 mL of LB+1.5% agar was pipetted onto the plate ∼1 h before imaging so that the agar could solidify completely. Once solid (after ∼5-10 min), a second Singer PlusPlate was placed on top of the agar pad to prevent contamination and drying. One microliter of cultures was spotted in the center of the pad and allowed to dry. A large 113 by 77 mm custom-made no. 1.5 glass coverslip (Nexterion) was applied [63]. Imaging was carried out in a heated environmental chamber with a water bubbler and several reservoirs of water to humidify the chamber. Phase images were acquired on a Nikon Ti-E inverted microscope every 5 min using a 40X air objective (NA: 0.95) with 1.5X magnification. Images were compiled into movies and analyzed using Matlab or FIJI. All rod-shaped cells were identified in the first frame and defined as able to grow if they at least doubled in mass and divided without lysis during the experiment.

### Fluorescent D-amino acid staining

HADA [64] was added to cultures at a final concentration of 500 μM during the last mass doubling of growth. To reduce background staining, cultures were diluted 1:10 in MSgg solution (5 mM potassium phosphate buffer + 0.05 M MOPS at pH 7) and then spotted onto a pad of that solution made with 1.5% agar. Cells were imaged using a Nikon Ti-E inverted microscope with a 100X oil objective (NA: 1.4). Phase and fluorescence (DAPI filter, ex/em: 375/460 nm, exposure time 2 s) images were acquired. Phase and fluorescence images were adjusted identically in Adobe Photoshop and merged in FIJI.

### Plating efficiency

Cultures were harvested and diluted 10-fold into LB. One hundred microliters of the dilutions were plated onto an LB plate. These plates were incubated overnight at 37 °C in a single layer (not stacked). CFU/mL values of the original culture were calculated from the colony counts of the dilutions that had distinct colonies.

### Lysis assay starting at different optical densities

Three colonies were inoculated into a single 10 mL LB culture and mixed well. Six two-fold serial dilutions of the culture were carried out and 5 mL of each dilution were transferred to a test tube and incubated at 37 °C on a roller drum until the undiluted culture reached an OD_600_∼0.8. Two hundred microliters of each culture were then aliquoted into a 96-well plate, the plate was sealed with optical film, and OD_600_ was monitored in a Biotek Epoch2 spectrophotometer.

### Oxygen depletion in an anaerobic chamber

Cultures were grown in aerobic conditions (96-well plate sealed with optical film, with holes poked through the film for each well). Cultures were then transferred into a Coy anaerobic chamber and OD_600_ was monitored using a Biotek Epoch2 spectrophotometer as above.

### Biophysical model linking oxygen diffusivity and concentration in a growing culture

Since the addition of surfactin increased oxygen levels in late-exponential phase of a growing culture (Fig. 3D), we sought to understand whether oxygen diffusivity could explain the increase. In a small region of extent Δ*l* at a depth *l* below the air-liquid interface of the culture, oxygen is depleted at a rate γ × *ρ*(*l*) × *c*(*l*)Δ*l*, where γ is the absorption rate per unit oxygen in close proximity to a cell, *ρ*(*l*) is the cell density at depth *l*, and *c*(*l*) is the oxygen concentration at depth *l*. At steady state, the rate of oxygen uptake in the region of extent Δ*l* must be balanced by the rate of supply via diffusion, that is

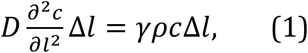

where *D* is the diffusivity of oxygen. The solution to Eq. 1 is 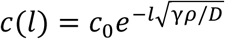, assuming that *c*_0_ is the oxygen concentration at the surface, *c* → 0 as *l* → ∞, and *γρ*/*D* is independent of *l*. Hence, as *D* increases, oxygen reaches increased depths as ∼ 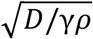, and the total amount of oxygen in the culture increases as 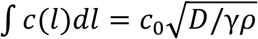. Thus, a 1.2-fold inc in the total amount of oxygen corresponds to a 1.4-fold increase in *D* of ∼ 1.4-fold, assuming that *c*_0_ is linked to solubility and *γ* and *ρ* remain approximately constant.

### Membrane potential measurements

Membrane potential was measured using a protocol modified from [42]. Cells were grown in 5 mL LB in a roller drum to OD_600_∼1. Cells were washed in a buffer containing 10 mM potassium phosphate, 5 mM MgSO4, and 250 mM sucrose (pH 7.0) and then resuspended to a calculated OD_600_ of 0.085 in that same buffer (pelleting steps were at 5100 rcf for 3 min). Two hundred microliters of this mixture were added to wells in a 96-well plate. 1 μM DiSC3(5) or 10 μM Thioflavin T (ThT) were added to the wells. Fluorescence readings (em/ex: 620/685 for DISC and 450/482 for ThT) were taken every 12 s with 5 s of linear shaking (567 cpm, 3-mm magnitude) on a Biotek Synergy H1 spectrophotometer. Readings were collected for ∼3 min before the drug was added and then readings were collected for another 10 min.

## Acknowledgements

The authors thank Alfred Spormann, Kyler Lugo, and the Huang lab for helpful discussions, Chao Jiang for help with strain construction, and Maya Farha and Eric Brown for help with membrane potential measurements. The authors acknowledge support from the Allen Discovery Center at Stanford on Systems Modeling of Infection (to H.A.A. and K.C.H.), National Institutes of Health (NIH) grants R35 GM131783 (to C.M.D. and D.B.K.) and R01 CA167250 (to C.A.D. and C.L.F.), and the Stanford Summer Research Amgen Scholars Program (to L.V.). K.C.H. is a Chan Zuckerberg Biohub Investigator.

## Supplemental Figures

**Figure S1:**
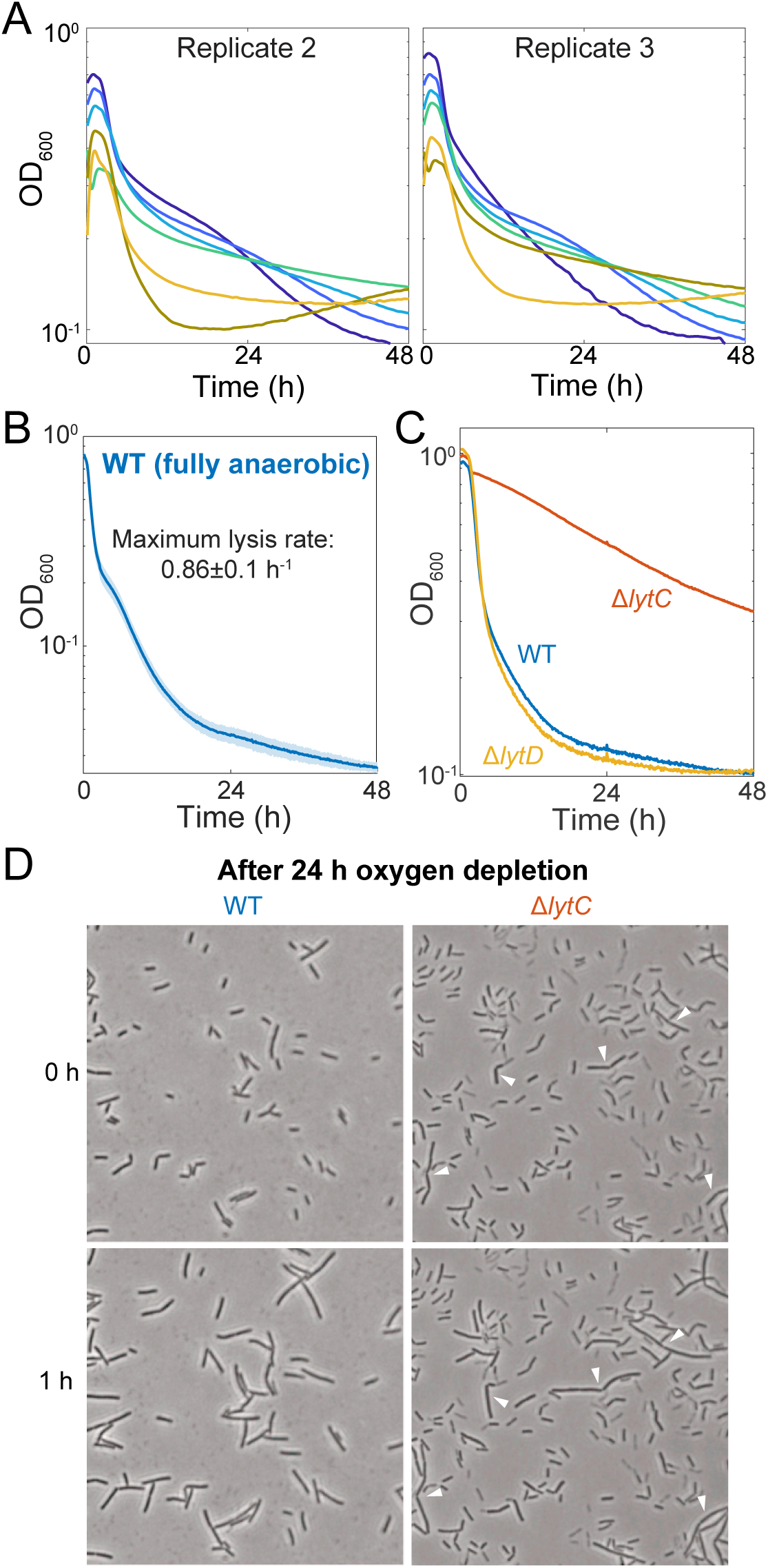
Characterization of depletion in an anaerobic chamber and of Δ*lytC* and Δ*lytD* mutants. (Related to Figure 1) (A) Culture lysis depends on cell density at the time of oxygen depletion. Two sets of independent lysis curves that vary in initial OD_600_ are shown; see Fig. 1D for the other independent experiments. (B) Cell lysis is more rapid under anaerobic conditions. Cultures were grown in 96-well plates to an OD_600_∼1 and then transferred to an anaerobic chamber to rapidly remove oxygen from the culture. Line represents the average and shading represents 1 SD, *n*=3. (C) Δ*lytD* cells show similar rates of lysis to wild-type (WT) 3610 cells upon oxygen depletion, unlike Δ*lytC* cells that exhibit slower lysis. Strains were grown to an OD600 ∼1 and then oxygen was depleted at *t* = 0. (D) After oxygen-depletion, the majority of WT cells but only a small subset of Δ*lytC* cells resume growth. After 24 h of oxygen depletion, WT and Δ*lytC* cultures were spotted onto an LB pad with oxygen and cell growth was monitored. Nearly all WT cells grew but only a few Δ*lytC* cells grew (arrowheads). Images shown are frames from Supplemental movies 1 and 2.

**Figure S2:**
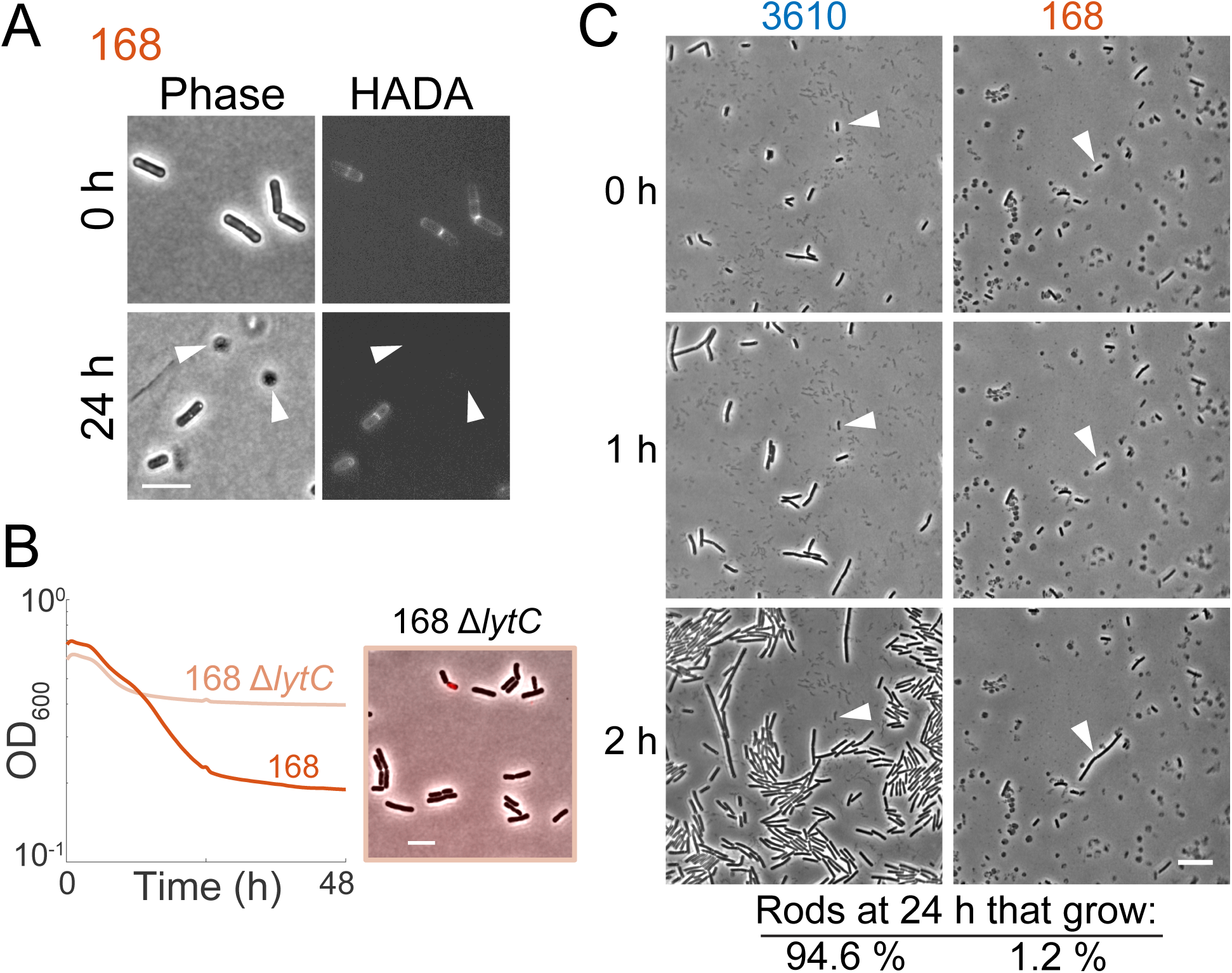
Oxygen depletion causes production of wall-less protoplasts in 168 and a low percentage of cells that are able to resume growth. (Related to Figure 2) (A) Cell-wall staining reveals that protoplasts have lost their cell wall. Phase-contrast and HADA (cell wall) staining of 168 cultures at 24 h post-oxygen depletion. Cells were stained with HADA for the last doubling before oxygen was depleted and the dye was present during the duration of depletion. Orange arrowheads: phase-dark spherical cells that lack cell wall staining. Scale bar: 5 μm. (B) 168 Δ*lytC* mutants remain rod-shaped following oxygen depletion. Left: OD_600_ of 168 Δ*lytC* exhibited somewhat slower lysis than the parent. Right: overlay of phase-contrast and propidium iodide staining of 168 Δ*lytC* cells. (C) Almost all 3610 cells grow once oxygen is restored following 24 h of oxygen depletion, but only ∼1% of 168 cells grow in similar circumstance. Cells from 3610 and 168 cultures after 24 h of oxygen depletion were spotted onto an LB pad and imaged in time-lapse to determine which cells were capable of growing and dividing. For 3610, the arrowhead points to a rare rod-shaped cell that did not grow. For 168, the arrowhead points to a rare rod-shaped cell that grew. Scale bar: 10 μm. *n* = 536 and 604 rod-shaped cells for 3610 and 168, respectively. Images shown are frames from Movies S3 and S4.

**Figure S3:**
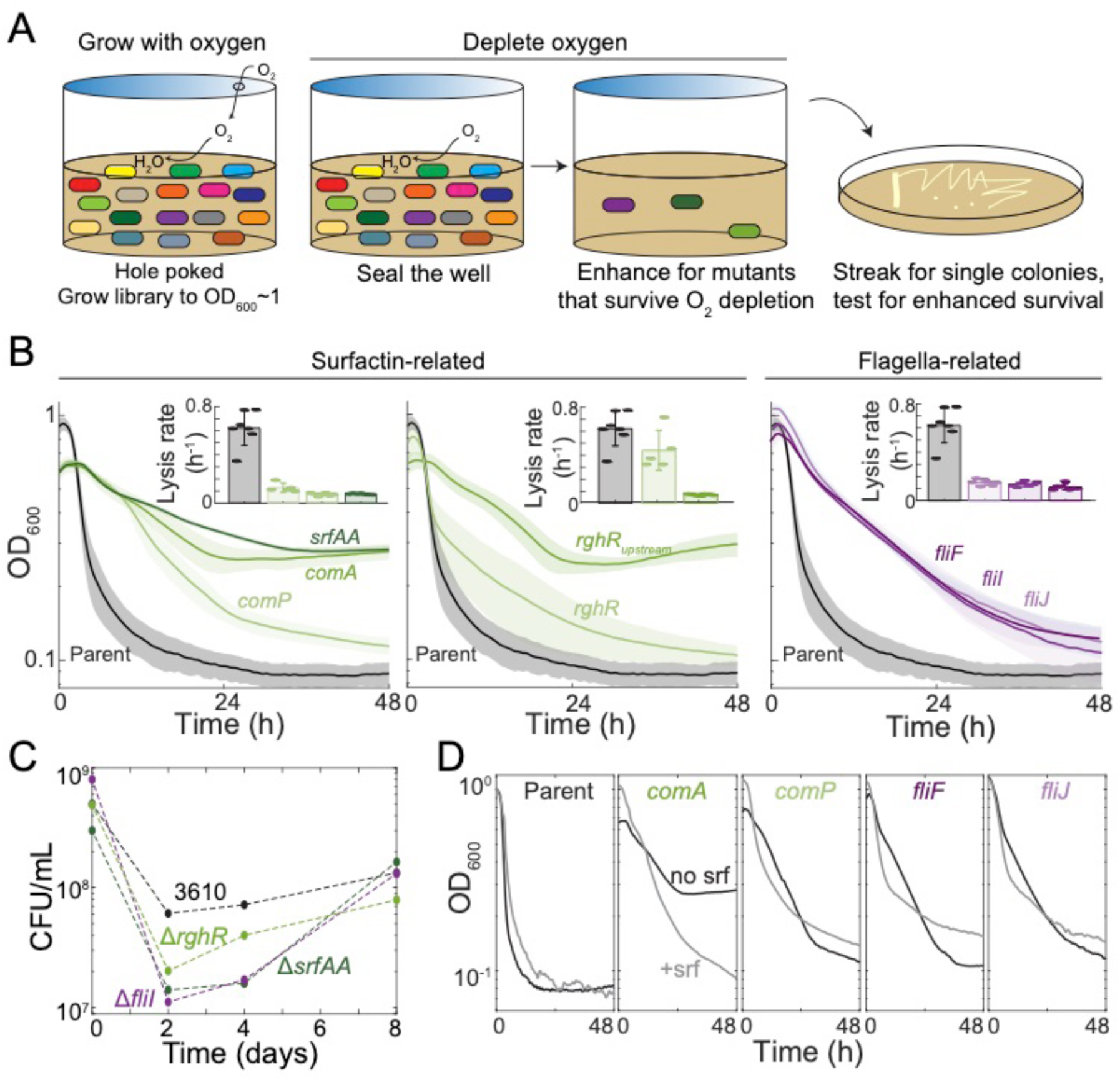
Characterization of hits from transposon screen. (Related to Figure 4) (A) Schematic of transposon screen design. A transposon mutant library was grown to an OD_600_∼1, and then oxygen was depleted. We screened cells that were able to recover after oxygen depletion for oxygen depletion phenotypes. (B) Transposon mutants have a reduced lysis rate. OD_600_ curves during oxygen depletion of cultures related to surfactin regulation (green) or flagella (purple). Lines represent the average and shading represents 1 SD, *n*=5. Inset: maximum lysis rates. Error bars represent 1 SD, *n*=5. (C) Plating efficiency of clean deletions of *srfAA, fliI*, and *rghR* are lower than that of 3610 2 and 4 days following oxygen depletion. (D) Addition of 48 µM surfactin increases the lysis rate of hits from transposon mutant screen.

**Figure S4:**
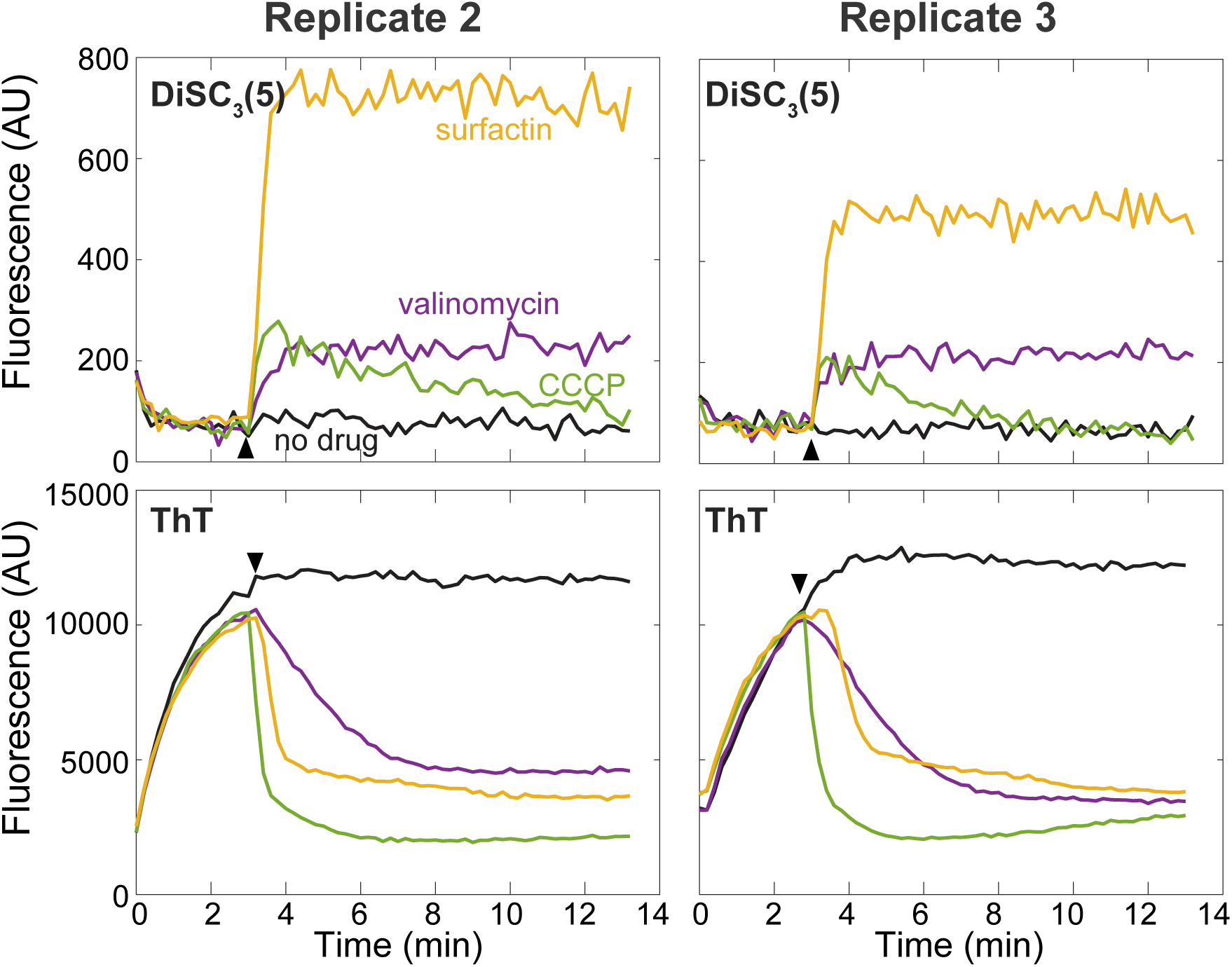
Membrane potential assays of 168 cells. (Related to Figure 5) DiSC_3_(5) (top) and ThT (bottom) both show that surfactin (48 μM), valinomycin (50 μM) and CCCP (5 μM) (time of addition marked by the black arrow) cause depolarization. Two experimental replicates are shown (replicate 1 is shown in Fig. 5).

## Supplemental Movie Legends

**Supplemental Movie 1:** The majority of oxygen-depleted cells from strain 3610 grow once oxygen is restored. Cells from a 24 h oxygen-depleted wild-type 3610 culture were spotted onto an LB pad and imaged every 5 min at 37 °C. This experiment was done in tandem with the 3610 Δ*lytC* cultures in Movie S2.

**Supplemental Movie 2:** A small subset of oxygen-depleted cells in the 3610 Δ*lytC* cultures grow once oxygen is restored. Cells from a 24 h oxygen-depleted Δ*lytC* culture were spotted on an LB pad and imaged every 5 min at 37 °C. This experiment was done in tandem with the 3610 cultures in Movie S1.

**Supplemental Movie 3:** The majority of oxygen-depleted cells from strain 3610 grow once oxygen is restored. This experiment was carried out as in Movie S1, and was done in tandem with the 168 cultures in Movie S4.

**Supplemental Movie 4:** A very small subset of oxygen-depleted, rod-shaped cells from strain 168 grow once oxygen is restored. Cells from a 24 h oxygen-depleted culture were spotted on an LB pad and imaged every 5 min at 37 °C. This experiment was done in tandem with the 3610 cultures in Movie S3.

## Supplemental Tables

**Table S1:**
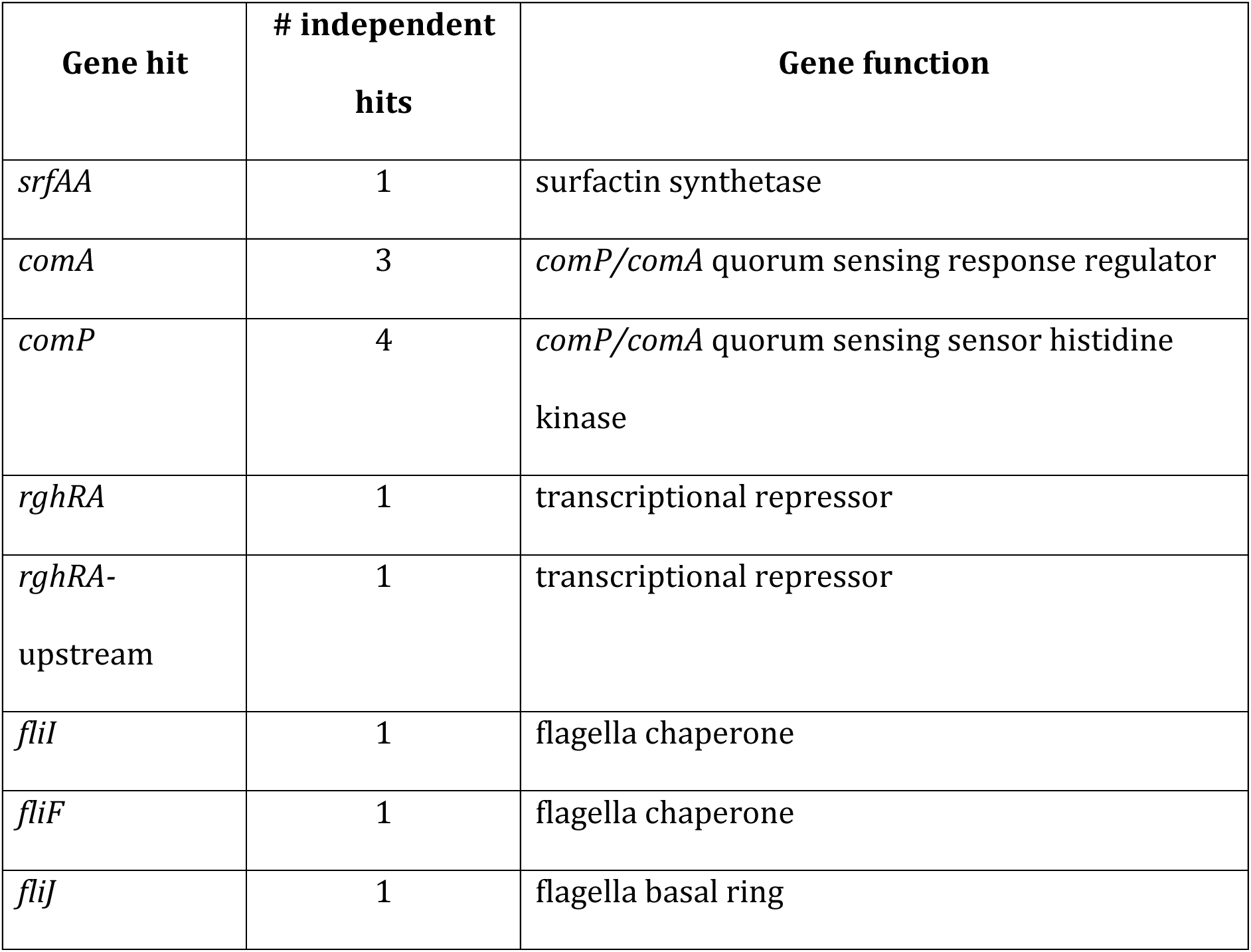
Information about hits from transposon screen.

**Table S2:**
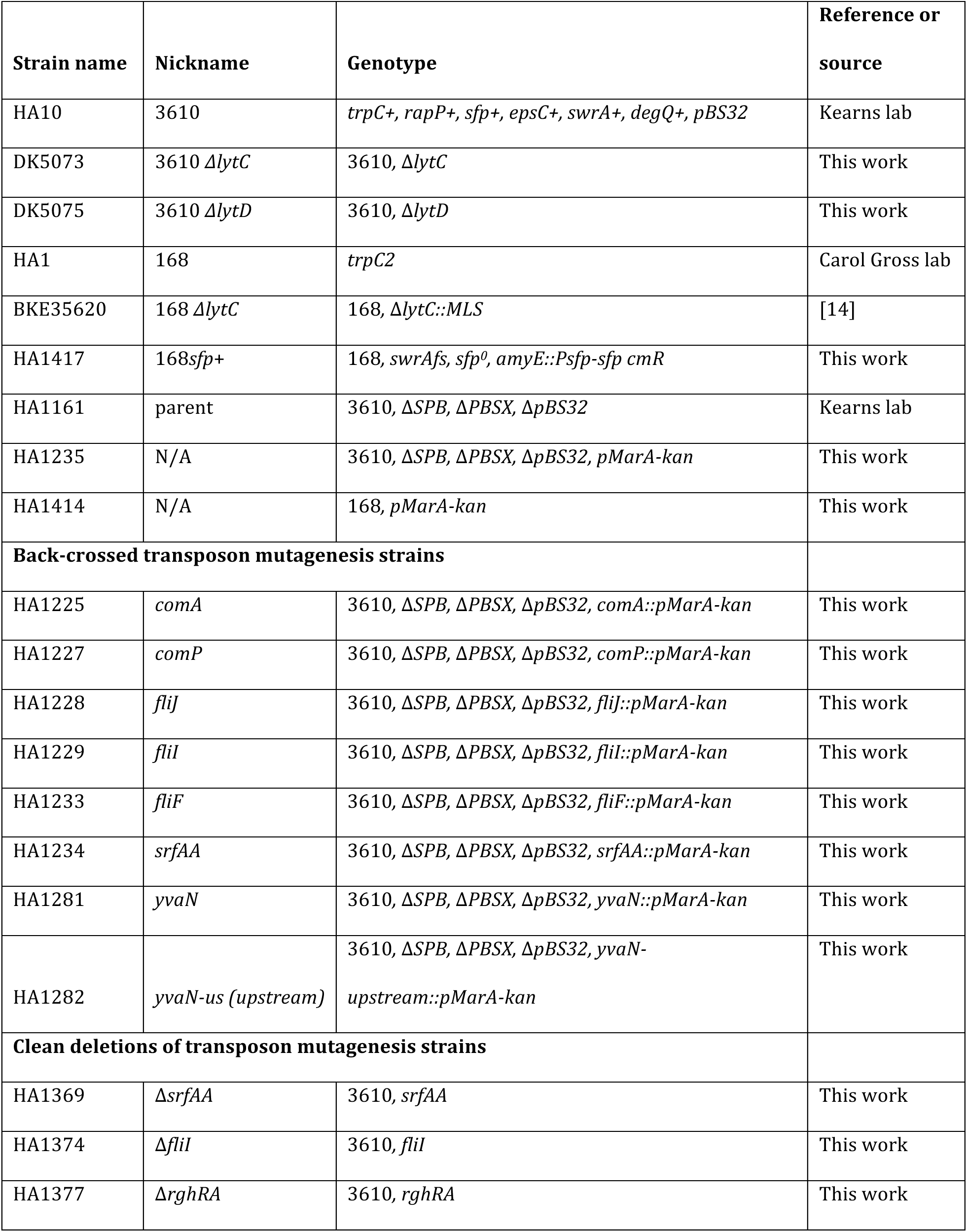
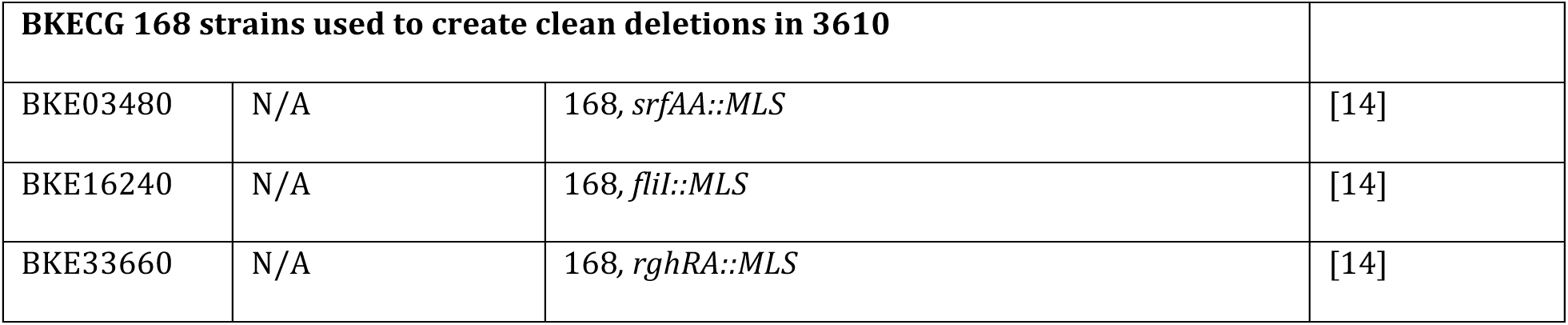
List of strains used in this study.

**Table S3:**
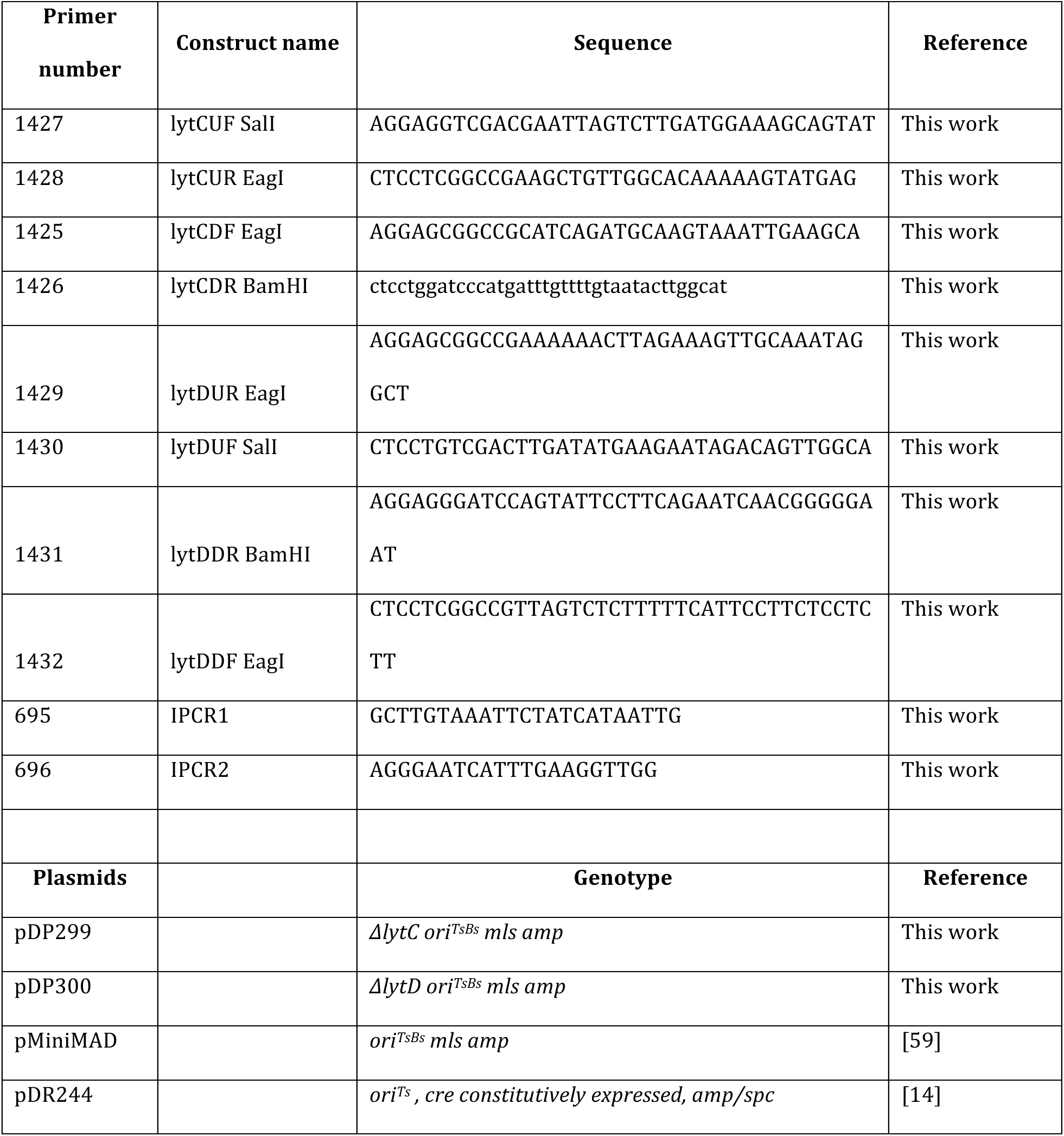
Primers and plasmids used in this study.

## References

1. Boutilier, R.G. (2001). Mechanisms of cell survival in hypoxia and hypothermia. J Exp Biol 204, 3171–3181.

2. Hochachka, P.W. (1986). Defense strategies against hypoxia and hypothermia. Science 231, 234–241.

3. Sendoel, A., and Hengartner, M.O. (2014). Apoptotic cell death under hypoxia. Physiology (Bethesda) 29, 168–176.

4. Prescott, L.M., Harley, J.P., and Klein, D.A. (2005). Microbiology, 6th Edition, (Dubuque, IA: McGraw-Hill Higher Education).

5. Wayne, L.G., and Hayes, L.G. (1996). An in vitro model for sequential study of shiftdown of *Mycobacterium tuberculosis* through two stages of nonreplicating persistence. Infect Immun 64, 2062–2069.

6. Rao, S.P., Alonso, S., Rand, L., Dick, T., and Pethe, K. (2008). The protonmotive force is required for maintaining ATP homeostasis and viability of hypoxic, nonreplicating *Mycobacterium tuberculosis*. Proc Natl Acad Sci U S A 105, 11945–11950.

7. Berney, M., Greening, C., Conrad, R., Jacobs, W.R., Jr., and Cook, G.M. (2014). An obligately aerobic soil bacterium activates fermentative hydrogen production to survive reductive stress during hypoxia. Proc Natl Acad Sci U S A 111, 11479–11484.

8. O’Toole, R., Smeulders, M.J., Blokpoel, M.C., Kay, E.J., Lougheed, K., and Williams, H.D. (2003). A two-component regulator of universal stress protein expression and adaptation to oxygen starvation in *Mycobacterium smegmatis*. J Bacteriol 185, 1543–1554.

9. Nakano, M.M., Dailly, Y.P., Zuber, P., and Clark, D.P. (1997). Characterization of anaerobic fermentative growth of *Bacillus subtilis*: identification of fermentation end products and genes required for growth. J Bacteriol 179, 6749–6755.

10. Pusey, P.L., and Wilson, C.L. (1984). Postharvest biological control of stone fruit brown rot by *Bacillus subtilis*. Plant Disease 68, 753–756.

11. Kumar, A.S., Lakshmanan, V., Caplan, J.L., Powell, D., Czymmek, K.J., Levia, D.F., and Bais, H.P. (2012). Rhizobacteria *Bacillus subtilis* restricts foliar pathogen entry through stomata. The Plant Journal 72, 694–706.

12. Sierra, J., and Renault, P. (1998). Temporal Pattern of Oxygen Concentration in a Hydromorphic Soil. Soil Science Society of America Journal 62, 1398–1405.

13. Kaufman, W., and Bauer, K. (1958). Some studies of the mechanism of the” anaerobic autolysis” of *Bacillus subtilis* J. Gen. Microbiol. 18.

14. Koo, B.M., Kritikos, G., Farelli, J.D., Todor, H., Tong, K., Kimsey, H., Wapinski, I., Galardini, M., Cabal, A., Peters, J.M., et al. (2017). Construction and Analysis of Two Genome-Scale Deletion Libraries for *Bacillus subtilis*. Cell Syst 4, 291–305 e297.

15. Peters, J.M., Colavin, A., Shi, H., Czarny, T.L., Larson, M.H., Wong, S., Hawkins, J.S., Lu, C.H.S., Koo, B.M., Marta, E., et al. (2016). A Comprehensive, CRISPR-based Functional Analysis of Essential Genes in Bacteria. Cell 165, 1493–1506.

16. Zhu, B., and Stulke, J. (2018). SubtiWiki in 2018: from genes and proteins to functional network annotation of the model organism *Bacillus subtilis*. Nucleic Acids Res 46, D743–D748.

17. Zeigler, D.R., Pragai, Z., Rodriguez, S., Chevreux, B., Muffler, A., Albert, T., Bai, R., Wyss, M., and Perkins, J.B. (2008). The origins of 168, W23, and other Bacillus subtilis legacy strains. J Bacteriol 190, 6983–6995.

18. Julkowska, D., Obuchowski, M., Holland, I.B., and Seror, S.J. (2005). Comparative analysis of the development of swarming communities of Bacillus subtilis 168 and a natural wild type: critical effects of surfactin and the composition of the medium. J Bacteriol 187, 65–76.

19. Nakano, M.M., Corbell, N., Besson, J., and Zuber, P. (1992). Isolation and characterization of sfp: a gene that functions in the production of the lipopeptide biosurfactant, surfactin, in Bacillus subtilis. Mol Gen Genet 232, 313–321.

20. Nakano, M.M., Marahiel, M.A., and Zuber, P. (1988). Identification of a genetic locus required for biosynthesis of the lipopeptide antibiotic surfactin in *Bacillus subtilis*. J Bacteriol 170, 5662–5668.

21. Arima, K., Kakinuma, A., and Tamura, G. (1968). Surfactin, a crystalline peptidelipid surfactant produced by *Bacillus subtilis*: isolation, characterization and its inhibition of fibrin clot formation. Biochem Biophys Res Commun 31, 488–494.

22. Płaza, G.A., Turek, A., Król, E., and Szczygłowska, R. (2013). Antifungal and antibacterial properties of surfactin isolated from *Bacillus subtilis* growing on molasses. African Journal of Microbiology Research 7, 3165–3170.

23. Korenblum, E., de Araujo, L.V., Guimaraes, C.R., de Souza, L.M., Sassaki, G., Abreu, F., Nitschke, M., Lins, U., Freire, D.M., Barreto-Bergter, E., et al. (2012). Purification and characterization of a surfactin-like molecule produced by Bacillus sp. H2O-1 and its antagonistic effect against sulfate reducing bacteria. BMC Microbiol 12, 252.

24. Rosenberg, G., Steinberg, N., Oppenheimer-Shaanan, Y., Olender, T., Doron, S., Ben-Ari, J., Sirota-Madi, A., Bloom-Ackermann, Z., and Kolodkin-Gal, I. (2016). Not so simple, not so subtle: the interspecies competition between *Bacillus simplex* and *Bacillus subtilis* and its impact on the evolution of biofilms. NPJ Biofilms Microbiomes 2, 15027.

25. Snook, M.E., Mitchell, T., Hinton, D.M., and Bacon, C.W. (2009). Isolation and characterization of leu7-surfactin from the endophytic bacterium *Bacillus mojavensis* RRC 101, a biocontrol agent for *Fusarium verticillioides*. J Agric Food Chem 57, 4287–4292.

26. Zhi, Y., Wu, Q., Du, H., and Xu, Y. (2016). Biocontrol of geosmin-producing Streptomyces spp. by two *Bacillus* strains from Chinese liquor. Int J Food Microbiol 231, 1–9.

27. Olmeda, B., Villen, L., Cruz, A., Orellana, G., and Perez-Gil, J. (2010). Pulmonary surfactant layers accelerate O(2) diffusion through the air-water interface. Biochim Biophys Acta 1798, 1281–1284.

28. Cabeen, M.T., and Jacobs-Wagner, C. (2005). Bacterial cell shape. Nat Rev Microbiol 3, 601–610.

29. Lazarevic, V., Margot, P., Soldo, B., and Karamata, D. (1992). Sequencing and analysis of the *Bacillus subtilis lytRABC* divergon: a regulatory unit encompassing the structural genes of the N-acetylmuramoyl-L-alanine amidase and its modifier. J Gen Microbiol 138, 1949–1961.

30. Margot, P., Mauel, C., and Karamata, D. (1994). The gene of the N-acetylglucosaminidase, a *Bacillus subtilis* 168 cell wall hydrolase not involved in vegetative cell autolysis. Mol Microbiol 12, 535–545.

31. Koch, A.L. (2001). Autolysis control hypotheses for tolerance to wall antibiotics. Antimicrob Agents Chemother 45, 2671–2675.

32. Sheppard, J.D., Jumarie, C., Cooper, D.G., and Laprade, R. (1991). Ionic channels induced by surfactin in planar lipid bilayer membranes. Biochim Biophys Acta 1064, 13–23.

33. Laidig, K.E., Gainer, J.L., and Daggett, V. (1998). Altering Diffusivity in Biological Solutions through Modification of Solution Structure and Dynamics. J Am Chem Soc 120, 9394–9395.

34. Hotez, L., Dailey, J.W., Geelhoed, G.W., and Gainer, J.L. (1977). The role of oxygen diffusivity in biochemical reactions. Experientia 33, 1424–1425.

35. Cosmina, P., Rodriguez, F., de Ferra, F., Grandi, G., Perego, M., Venema, G., and van Sinderen, D. (1993). Sequence and analysis of the genetic locus responsible for surfactin synthesis in *Bacillus subtilis*. Mol Microbiol 8, 821–831.

36. Nakano, M.M., Xia, L.A., and Zuber, P. (1991). Transcription initiation region of the *srfA* operon, which is controlled by the *comP-comA* signal transduction system in *Bacillus subtilis*. J Bacteriol 173, 5487–5493.

37. Hayashi, K., Kensuke, T., Kobayashi, K., Ogasawara, N., and Ogura, M. (2006). *Bacillus subtilis* RghR (YvaN) represses *rapG* and *rapH*, which encode inhibitors of expression of the *srfA* operon. Mol Microbiol 59, 1714–1729.

38. Mukherjee, S., and Kearns, D.B. (2014). The structure and regulation of flagella in *Bacillus subtilis*. Annu Rev Genet 48, 319–340.

39. Ueno, T., Oosawa, K., and Aizawa, S. (1992). M ring, S ring and proximal rod of the flagellar basal body of *Salmonella typhimurium* are composed of subunits of a single protein, FliF. J Mol Biol 227, 672–677.

40. Evans, L.D., Stafford, G.P., Ahmed, S., Fraser, G.M., and Hughes, C. (2006). An escort mechanism for cycling of export chaperones during flagellum assembly. Proc Natl Acad Sci U S A 103, 17474–17479.

41. Minamino, T., Chu, R., Yamaguchi, S., and Macnab, R.M. (2000). Role of FliJ in flagellar protein export in *Salmonella*. J Bacteriol 182, 4207–4215.

42. Farha, M.A., Verschoor, C.P., Bowdish, D., and Brown, E.D. (2013). Collapsing the proton motive force to identify synergistic combinations against *Staphylococcus aureus*. Chem Biol 20, 1168–1178.

43. Prindle, A., Liu, J., Asally, M., Ly, S., Garcia-Ojalvo, J., and Suel, G.M. (2015). Ion channels enable electrical communication in bacterial communities. Nature 527, 59–63.

44. Varma, S., Sabo, D., and Rempe, S.B. (2008). K+/Na+ selectivity in K channels and valinomycin: over-coordination versus cavity-size constraints. J Mol Biol 376, 13–22.

45. Strahl, H., Burmann, F., and Hamoen, L.W. (2014). The actin homologue MreB organizes the bacterial cell membrane. Nat Commun 5, 3442.

46. Branda, S.S., Gonzalez-Pastor, J.E., Ben-Yehuda, S., Losick, R., and Kolter, R. (2001). Fruiting body formation by *Bacillus subtilis*. Proc Natl Acad Sci U S A 98, 11621–11626.

47. Pham, J.V., Yilma, M.A., Feliz, A., Majid, M.T., Maffetone, N., Walker, J.R., Kim, E., Cho, H.J., Reynolds, J.M., Song, M.C., et al. (2019). A Review of the Microbial Production of Bioactive Natural Products and Biologics. Front Microbiol 10, 1404.

48. Hu, J., Lei, P., Mohsin, A., Liu, X., Huang, M., Li, L., Hu, J., Hang, H., Zhuang, Y., and Guo, M. (2017). Mixomics analysis of Bacillus subtilis: effect of oxygen availability on riboflavin production. Microb Cell Fact 16, 150.

49. Garcia-Ochoa, F., and Gomez, E. (2009). Bioreactor scale-up and oxygen transfer rate in microbial processes: an overview. Biotechnol Adv 27, 153–176.

50. Baeshen, N.A., Baeshen, M.N., Sheikh, A., Bora, R.S., Ahmed, M.M., Ramadan, H.A., Saini, K.S., and Redwan, E.M. (2014). Cell factories for insulin production. Microb Cell Fact 13, 141.

51. Arjes, H.A., Kriel, A., Sorto, N.A., Shaw, J.T., Wang, J.D., and Levin, P.A. (2014). Failsafe mechanisms couple division and DNA replication in bacteria. Curr Biol 24, 2149–2155.

52. Rojas, E.R., Huang, K.C., and Theriot, J.A. (2017). Homeostatic Cell Growth Is Accomplished Mechanically through Membrane Tension Inhibition of Cell-Wall Synthesis. Cell Syst 5, 578–590 e576.

53. Blackman, S.A., Smith, T.J., and Foster, S.J. (1998). The role of autolysins during vegetative growth of *Bacillus subtilis* 168. Microbiology 144 ( *Pt 1*), 73–82.

54. Calamita, H.G., Ehringer, W.D., Koch, A.L., and Doyle, R.J. (2001). Evidence that the cell wall of Bacillus subtilis is protonated during respiration. Proc Natl Acad Sci U S A 98, 15260–15263.

55. Mueller, E.A., Egan, A.J., Breukink, E., Vollmer, W., and Levin, P.A. (2019). Plasticity of *Escherichia coli* cell wall metabolism promotes fitness and antibiotic resistance across environmental conditions. Elife 8.

56. Peters, K., Kannan, S., Rao, V.A., Biboy, J., Vollmer, D., Erickson, S.W., Lewis, R.J., Young, K.D., and Vollmer, W. (2016). The Redundancy of Peptidoglycan Carboxypeptidases Ensures Robust Cell Shape Maintenance in *Escherichia coli*. MBio 7.

57. Stratford, J.P., Edwards, C.L.A., Ghanshyam, M.J., Malyshev, D., Delise, M.A., Hayashi, Y., and Asally, M. (2019). Electrically induced bacterial membrane-potential dynamics correspond to cellular proliferation capacity. Proc Natl Acad Sci U S A 116, 9552–9557.

58. Yasbin, R.E., and Young, F.E. (1974). Transduction in *Bacillus subtilis* by bacteriophage SPP1. J Virol 14, 1343–1348.

59. Patrick, J.E., and Kearns, D.B. (2008). MinJ (YvjD) is a topological determinant of cell division in *Bacillus subtilis*. Mol Microbiol 70, 1166–1179.

60. Arnaud, M., Chastanet, A., and Debarbouille, M. (2004). New vector for efficient allelic replacement in naturally nontransformable, low-GC-content, gram-positive bacteria. Appl Environ Microbiol 70, 6887–6891.

61. DeRosa, C.A., Samonina-Kosicka, J., Fan, Z., Hendargo, H.C., Weitzel, D.H., Palmer, G.M., and Fraser, C.L. (2015). Oxygen Sensing Difluoroboron Dinaphthoylmethane Polylactide. Macromolecules 48, 2967–2977.

62. DeRosa, C.A., Seaman, S.A., Mathew, A.S., Gorick, C.M., Fan, Z., Demas, J.N., Peirce, S.M., and Fraser, C.L. (2016). Oxygen Sensing Difluoroboron beta-Diketonate Polylactide Materials with Tunable Dynamic Ranges for Wound Imaging. ACS Sens 1, 1366–1373.

63. Shi, H., Colavin, A., Lee, T.K., and Huang, K.C. (2017). Strain Library Imaging Protocol: high-throughput, automated single-cell microscopy for large bacterial collections arrayed on multiwell plates. Nature Protocols.

64. Kuru, E., Hughes, H.V., Brown, P.J., Hall, E., Tekkam, S., Cava, F., de Pedro, M.A., Brun, Y.V., and VanNieuwenhze, M.S. (2012). In Situ probing of newly synthesized peptidoglycan in live bacteria with fluorescent D-amino acids. Angew Chem Int Ed Engl 51, 12519–12523.

